# CO_2_-sensitive Cx26 hemichannels in dorsal raphe-ventral tegmental area pathway mediate hypercapnic arousal

**DOI:** 10.1101/2025.04.14.648741

**Authors:** Lumei Huang, Jack Butler, Alexandra Lovatt, Amol Bhandare, Julien Pelletier, Julia Baran, Emily Lane-Hill, Nicholas Dale, Mark J. Wall

## Abstract

Hypercapnic arousal is a life-preserving reflex that promotes awakening in response to elevated PCO_2_, yet the cellular mechanisms by which CO_2_ is detected remain incompletely understood. Serotonergic neurons of the dorsal raphe (DR) have been implicated in hypercapnic responses, but the molecular sensor that directly detects CO_2_ within this circuit is unknown. Here, we show that the CO_2_-gated connexin 26 (Cx26) hemichannel is expressed on serotonergic terminals projecting from the DR to the ventral tegmental area (VTA). Brief elevations in PCO_2_ modulated the excitability of VTA dopamine neurons through serotonergic signaling without altering glutamatergic transmission, and these effects were abolished by pharmacological inhibition of hemichannels. By expressing Cx26 and genetically encoded GRAB_5HT_ sensors in cultured cells, we further demonstrate that Cx26 hemichannels opened by elevated PCO_2_ are directly permeable to serotonin. Selective removal of CO_2_ sensitivity from Cx26 in DR serotonergic neurons delayed arousal from sleep during transient hypercapnia and reduced activation of VTA dopamine neurons *in vivo*. Together, these findings identify Cx26 hemichannels as a direct CO_2_ sensor that enables serotonin release through a channel co-synapse and contributes to hypercapnic arousal.

## INTRODUCTION

Hypercapnia, excessive CO_2_ retention (>∼45 mm Hg PCO_2_), frequently occurs in patients with obstructive sleep apnoea (OSA) (*1*) or with chronic obstructive pulmonary disease (COPD) (*2*). The immediate activation of arousal-promoting brain circuitry by hypercapnia is an important lifesaving chemoreflex that rapidly re-establishes upper airway patency and restores ventilation (*3, 4*).

Several brain regions contribute to hypercapnia-induced arousal (*5*). These include glutamatergic neurons and calcitonin gene-related peptide (PBel ^CGRP^ neurons) in the lateral parabrachial nucleus (PB) (*6-8*), noradrenergic neurons in the locus coeruleus (LC) (*9, 10*), histaminergic neurons in the tuberomammillary nucleus (TMN) (*11, 12*), orexinergic neurons in the lateral hypothalamus (LH) (*6*), parvalbumin (PV) neurons in basal forebrain (*13, 14*), and serotoninergic neurons (5-HT) in the dorsal raphe (DR) (*8, 15, 16*). Genetic and optogenetic manipulations have shown that 5-HT neurons from DR projecting to PBel ^CGRP^ neurons play an important role in hypercapnic arousal (*8*). The main mechanism of circuit activation to produce hypercapnia-induced wakefulness is conventionally attributed to changes in pH, resulting from the conversion of CO_2_ into an anion (HCO^3-^) and a hydrogen ion (H^+^), rather than by direct CO_2_ sensing. Dopaminergic (DA) neurons in the ventral tegmental area (VTA) play a fundamental role in the maintenance of the wakefulness state (*17, 18*). Recently, two additional VTA neuronal populations, GABAergic neurons and glutamatergic neurons have been reported to either inhibit or promote wakefulness, respectively (*19*). Apart from local circuitry, VTA neurons also receive strong upstream projections from other sleep/arousal-mediating brain regions, including DR (*8, 20*), LH (*21, 22*), pedunculopontine tegmental nucleus (PPTg) and laterodorsal tegmental nucleus (LDTg) (*23*). The innervation of VTA from 5-HT neurons in DR directly regulates wakefulness (*20, 24*). However, this DR-VTA pathway has not previously been implicated in hypercapnic arousal.

Connexin 26 (Cx26) hemichannels directly sense CO_2_ (*25-28*) and are fully opened when PCO_2_ exceeds ∼55 mmHg (hypercapnia). Although Cx26 has been considered a glial cell connexin (*29, 30*), neuronal Cx26 expression can be detected in substantia nigra pars compacta dopaminergic neurons (*31*), in subplate zone neurons (*32, 33*), and in VTA GABAergic neurons (*34*).

Electrophysiological recording confirmed that the Cx26 hemichannels expressed by VTA GABAergic neurons directly regulate their excitability in response to rising levels of PCO_2_ (*34*). However, it is unclear whether Cx26 hemichannels are also expressed by other VTA neurons to further modulate neural circuits during hypercapnia.

We investigated whether direct CO_2_ sensing mediated by Cx26 hemichannels contributes to hypercapnic arousal through the DR–VTA circuit. Specifically, we examined whether Cx26 is expressed in DR serotonergic neurons and their projections to the VTA, whether elevated PCO_2_ triggers serotonin release through Cx26 hemichannels, and whether this signaling pathway influences VTA neuronal activity and behavioral arousal. These experiments address how direct CO_2_ detection is coupled to neural circuits that promote awakening during hypercapnia.

## RESULTS

### Cx26 hemichannels are expressed in the VTA

Following our previous discovery that ventral tegmental area (VTA) GABAergic neurons express Cx26 hemichannels that are sensitive to CO_2_ (*34*), we sought to map the functional expression of CO_2_-sensitive Cx26 hemichannels within VTA. we adapted a well-established dye-loading assay in which the membrane-impermeant fluorescent dye fluorescein 5(6)-isothiocyanate (FITC) enters cells through open Cx26 hemichannels under conditions of elevated PCO_2_ at constant extracellular pH(*25, 34, 35*). To label CO_2_-sensitive structures in vivo, we combined this assay with transcardial perfusion (**see Methods**). Mice were perfused with a 55 mmHg PCO_2_ aCSF solution containing FITC (50 μM) for 3 min (2.5–3 ml min^-1^), followed by perfusion with 20 mmHg PCO_2_ aCSF without FITC for 3 min. Control animals received 20 mmHg PCO_2_ aCSF containing FITC for 3 min followed by the same level of CO_2_ without FITC for 3 min (**Fig. 1A**). Robust FITC labelling of blood vessels (including endothelial cells) and astrocytes, which are known to express functional Cx26 hemichannels, confirmed the success of this brief transcardial dye-loading protocol (**Fig. S 1A-B**).

**Fig. 1.**
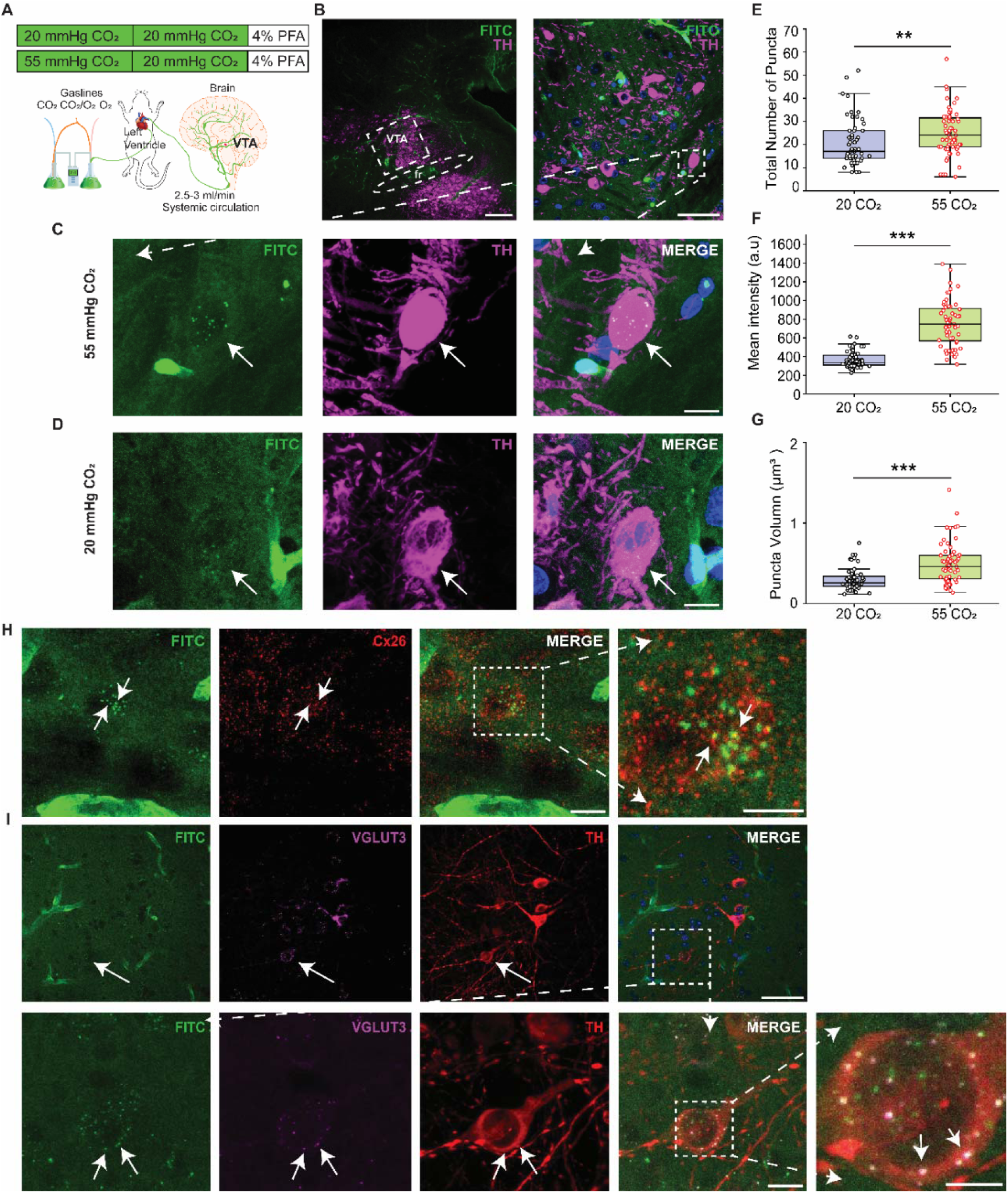
Elevated PCO_2_ loads Cx26_2_ and VGLUT3_2_ puncta surrounding VTA DA neurons. **(A)** Schematic of the transcardial perfusion protocol using FITC (green) dissolved in aCSF equilibrated with either 20 or 55 mmHg CO_2_ to enable dye loading through CO_2_-sensitive channels in the brain. (**B)** Confocal images showing FITC fluorescence and tyrosine hydroxylase (TH) staining of the VTA in the sagittal plane (left panel scale bar 200 µm; right panel scale bar 50 µm). (**C and D)** High magnification images showing FITC-loaded puncta (green) and TH-positive neurons (magenta) in 20 mmHg CO_2_ and 55 mmHg CO_2_ groups (scale bar 10 µm). (**E-G)** Quantification of the FITC-loaded puncta surrounding TH-positive neurons (20 mmHg CO_2_ only: n = 65 cells; 55 mmHg CO_2_: n = 55 cells). In comparison with the control group (20 mmHg CO_2_ only), perfusion with 55 mmHg CO_2_ significantly increased puncta number **(E)**, fluorescent intensity **(F)**, and puncta volume **(G).** (**H-I)** Images showing the colocalization of FITC-loaded puncta with Cx26 and VGLUT3, respectively (scale bar 10 µm, insert images, scale bar 2 µm). Data are mean ± SEM. Statistical significance was assessed using Mann–Whitney tests. ***P < 0.0001, **P < 0.002.

Although this brief elevation of PCO_2_ did not result in FITC loading of neuronal somata, numerous FITC-positive puncta were observed throughout the VTA. Interestingly, these puncta were clustered around NeuN^+^ neurons (**Fig. S 1C**). To further identify the neuron types, we used the DA neuron marker tyrosine hydroxylase (TH) and found that the FITC-loaded puncta surrounded DA neurons (**Fig. 1B-D**). Compared with control animals perfused at 20 mmHg PCO_2_, perfusion with 55 mmHg PCO_2_ significantly increased the number of FITC-positive puncta surrounding DA neurons (**Fig. 1E**), as well as their fluorescence intensity (**Fig. 1F**) and volume (**Fig. 1G**). FITC-positive puncta were also observed around non-DA neurons in the VTA, although these populations were not further characterized **(Fig. 1D)**. Many FITC-loaded puncta were Cx26^+^ (**Fig. 1H**), suggesting that dye entry through CO_2_-sensitive Cx26 hemichannels. The specificity of the antibody for Cx26 was confirmed in cultured cells (**Fig. S2A-E**). To determine whether these puncta expressed in presynaptic terminals, sections were stained for the vesicular glutamate transporter VGLUT3. We found that FITC-loaded puncta colocalized with VGLUT3 (**Fig. 1I)** and some puncta also colocalized with VGLUT2 (**Fig. S1E)**. Together, these findings demonstrate that functional CO_2_-sensitive Cx26 hemichannels are expressed on presynaptic terminals surrounding VTA dopaminergic neurons within the VTA.

### Cx26 localizes to DR–VTA serotonergic terminals

To our knowledge, no neurons within VTA express VGLUT3, therefore VGLUT3-positive terminals identified in the VTA are likely to originate from afferent projections. A strong candidate is the DR, which provides serotonergic input to the VTA from neurons that co-express VGLUT3 and can release both glutamate and 5-HT (*36-38*). We therefore investigated whether Cx26 is expressed on serotonergic terminals within the VTA. Serotonergic axons and terminals can be identified by immunolabelling for the serotonin transporter (SERT) (*39*). We detected that Cx26 and VGLUT3 colocalized on SERT-positive fibres within the VTA (**Fig. 2A**). Furthermore, Cx26 labelled puncta together with SERT-positive puncta were found densely clustered around TH^+^-positive DA neurons (**Fig. 2B**), suggesting that Cx26 is expressed on serotonergic inputs to VTA DA neurons. The expression of Cx26 on VTA DA neurons was excluded, as we did not detect Cx26 mRNA (leptomeninges as a positive control for GJB2 expression **Fig. S3A–C** and **Fig. 4A–B)** in the VTA DA neurons. Instead, the expression of VTA Cx26 mRNA is restricted to specific GABAergic and VGLUT2-positive neuronal populations **(Fig. S4C–D)**.To further characterize the potential for CO_2_-dependent regulation of these terminals, we examined the expression of aquaporin 5 (AQP5), a neuronal aquaporin capable of facilitating CO_2_ permeation across cell membranes (*40, 41*). AQP5 was detected on SERT-positive fibres in the VTA (**Fig. 2C**), providing additional support to our hypothesis of CO_2_-dependent gating of Cx26 hemichannels on nerve terminals within the VTA.

**Fig. 2.**
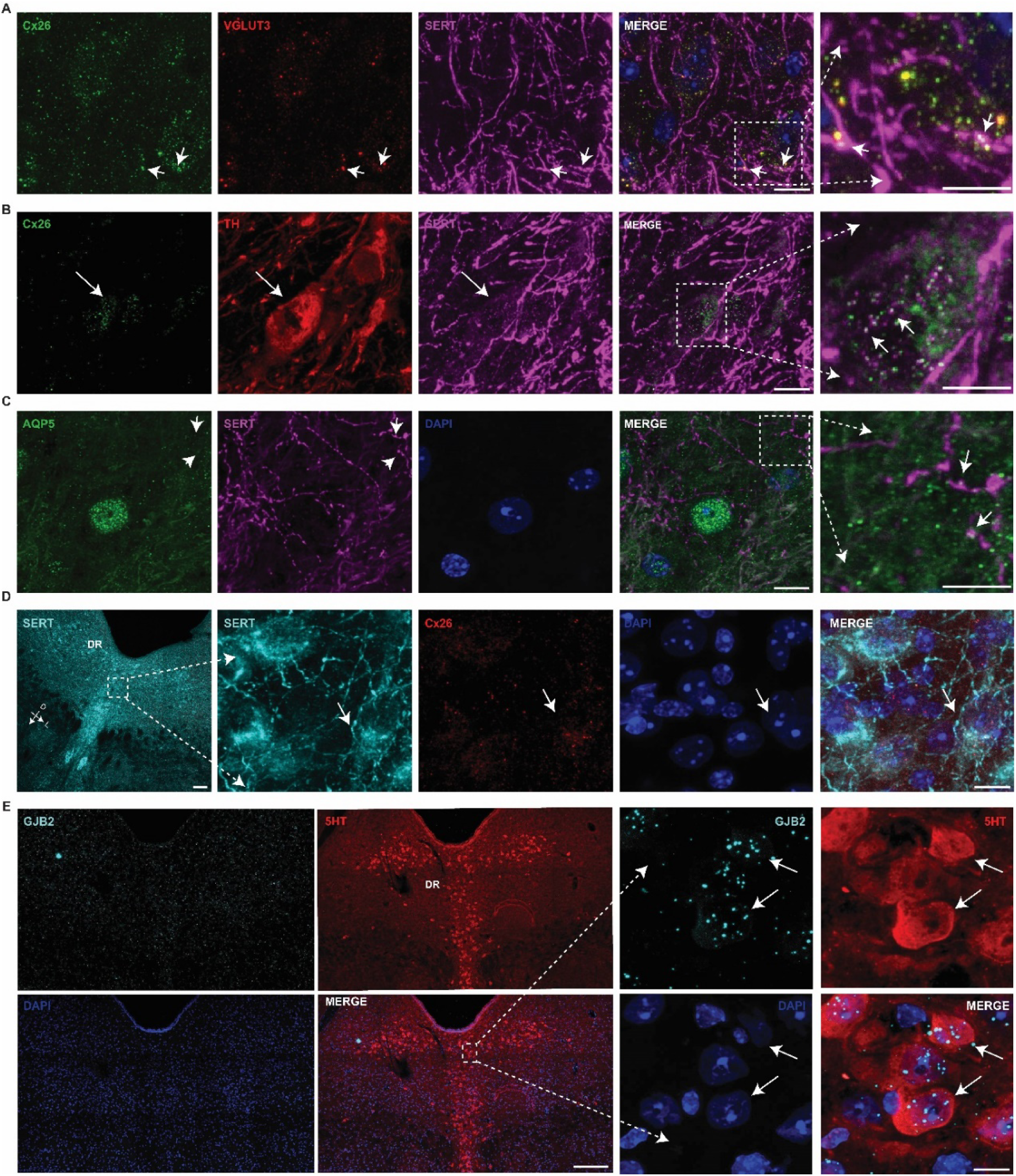
Connexin 26 is expressed in SERT-positive fibres in the VTA. **(A)** Confocal images of the VTA illustrating the expression of Cx26 (green) with VGLUT3 (red) on SERT-positive terminals (magenta). (**B)** Images illustrating that Cx26 (green) are localized on SERT-positive terminals (magenta) surround TH-positive neurons (red). (**C)** Images showing the localization of CO_2_-permeable aquaporin (AQP5: green) on SERT-positive fibres (magenta). (**D)** Images showing the somatic expression of Cx26 (red) on SERT-positive neurons in the DR. **E)** Confocal images of RNAscope labelling (*GJB2*: magenta) in combination with immunofluorescence (5-HT: green) showing *GJB2* mRNA (magenta) is located in 5-HT positive cells in the DR. Images from **A-D** (scale bar 10 µm; Dash square images scale bar 2 µm), Images from **E** (scale bar 200 µm; Dash square images scale bar 10 µm). The white arrows indicate colocalization. Each staining experiment was repeated three times.

In addition, our immunostaining (**Fig. 2D**) and RNAscope results revealed that DR 5-HT^+^ neurons express Cx26 (**Fig. 2E**). Elevated PCO_2_ increased the firing rate of DR neurons **(Fig. S10)**. Together, these findings identify the DR–VTA pathway as a major site of functional Cx26 hemichannels expression.

### Elevated PCO_2_ does not alter VTA neurotransmission

The expression of Cx26 on VGLUT3-positive serotonergic nerve terminals from DR indicates that the opening of Cx26 hemichannels could modulate glutamatergic or serotonergic transmission onto VTA DA neurons. For example, it is possible that opening of the hemichannels by raised PCO_2_ could enhance glutamatergic transmission by allowing Ca^2+^ influx into the nerve terminals. Therefore, we recorded miniature postsynaptic currents (mPSCs) in VTA DA neurons in the presence of 1 µM tetrodotoxin (TTX) to block action potentials. Recorded neurons were identified by their relative position in the slice, their electrophysiological properties (**Fig. 3A**), hyperpolarisation induced by dopamine (100 µM) (**Fig. 3B**) and post hoc TH staining of biocytin filled neurons (**Fig. 3F**). After 5 minutes of baseline recording in a physiological level of PCO_2_ (∼35 mmHg PCO_2_), slices were perfused with 55 mmHg PCO_2_ for 10 minutes, followed by 35 mmHg PCO_2_ to recover (**Fig. 3C**). Increasing PCO_2_ had no significant effect on either the inter-event interval or amplitude of the mPSCs (**Fig. 3D**). It is possible that hemichannel opening had distinct effects on different synapses (for example enhancing glutamatergic transmission but reducing GABAergic transmission), and thus the overall effect on the mPSCs was not significant. To measure glutamatergic transmission, we recorded AMPA receptor-mediated mEPSCs (in 1 µM TTX, 50 mM picrotoxin to block GABA_A_ receptors and 5 mM L689,560 to block NMDA receptors). Again, there were no significant differences in either the inter-event interval or amplitude of AMPA receptor-mediated mEPSCs (**Fig. 3E**), indicating that 55 mmHg CO_2_ did not significantly change glutamatergic transmission onto VTA DA neurons.

**Fig. 3.**
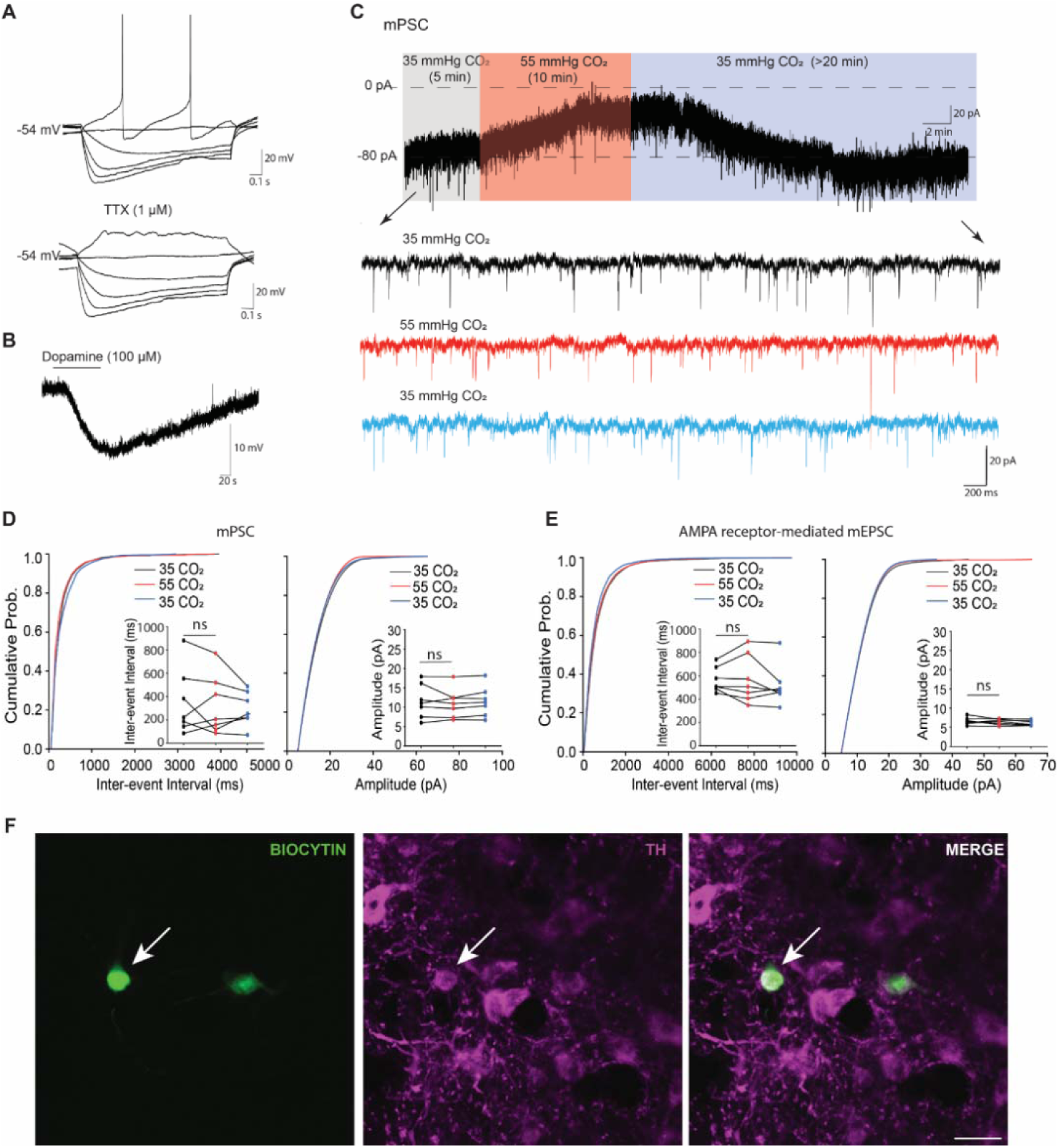
Increases in PCO2 do not change spontaneous synaptic currents recorded in VTA DA neurons. **(A)** Representative current–voltage relationship of a VTA DA neuron (upper panel). Application of tetrodotoxin (TTX; 1 μM) abolished spontaneous action potential firing (lower panel). (**B)** Application of dopamine (100 µM) hyperpolarized the membrane potential. (**C)** Representative voltage-clamp recordings (holding potential, −70 mV) of miniature postsynaptic currents (mPSCs) obtained under control (35 mmHg CO_2_) and hypercapnic (55 mmHg CO_2_ for 10 min) conditions. The upper panel shows the full recording, while the lower panels display expanded segments of mPSCs. Note the change in holding current observed during exposure to 55 mmHg CO_2_. (**D and E)** Cumulative probability distributions of mPSC (**D**, *n* = 7 cells) and AMPA receptor-mediated miniature excitatory postsynaptic current (mEPSC; **E**, *n* = 6 cells) inter-event intervals and amplitudes. AMPA receptor-mediated mEPSCs were isolated by pharmacological blockade of GABAA and NMDA receptors. Insets show mean inter-event intervals and amplitudes. Elevation of PCO_2_ did not significantly affect either the frequency (inter-event interval) or amplitude of mPSCs. Likewise, no significant changes were observed in the frequency or amplitude of AMPA receptor-mediated mEPSCs. **(F)** Post hoc identification of recorded DA neurons. Representative images show biocytin-filled neurons colocalized with the dopaminergic marker tyrosine hydroxylase (TH, magenta). Scale bar: 50 μm. Data are mean ± SEM. Statistical significance was assessed using one-way repeated-measures ANOVA followed by Tukey’s multiple-comparisons test. ns, not significant.

### Elevated PCO_2_ changes VTA DA firing via 5-HT receptors

Although no observable changes in glutamatergic transmission onto VTA DA neurons were detected when PCO_2_ was increased, clear changes in the holding current were observed (**Fig. 3C**), suggesting that membrane properties and potentially neuronal excitability may be affected. Therefore, we switched to current clamp recording and directly examined the effects of elevated PCO_2_ on spontaneous action potential firing. Following baseline recordings in control aCSF (∼35 mmHg PCO_2_), slices were exposed to 55 mmHg PCO_2_ for 1 min. Out of 29 DA neurons recorded, 17 neurons (∼59 %) increased firing rate by ∼37% in response to elevated PCO_2_ (**Fig. 4B and H**), whereas the remaining 12 neurons (∼41%) exhibited a decrease in firing rate of approximately 52% (**Fig. 4C and I**). These changes were not caused by mechanical disturbance during solution exchange, as no changes in firing rate were observed (**Fig. 4A)** when perfusion was switched between two bottles containing the same aCSF equilibrated with 35 mmHg CO_2_. Plotting firing rates during elevated PCO_2_ against baseline firing rates clearly separated neurons exhibiting excitatory and inhibitory responses from the control group (**Fig. 4M Left**).

**Fig. 4.**
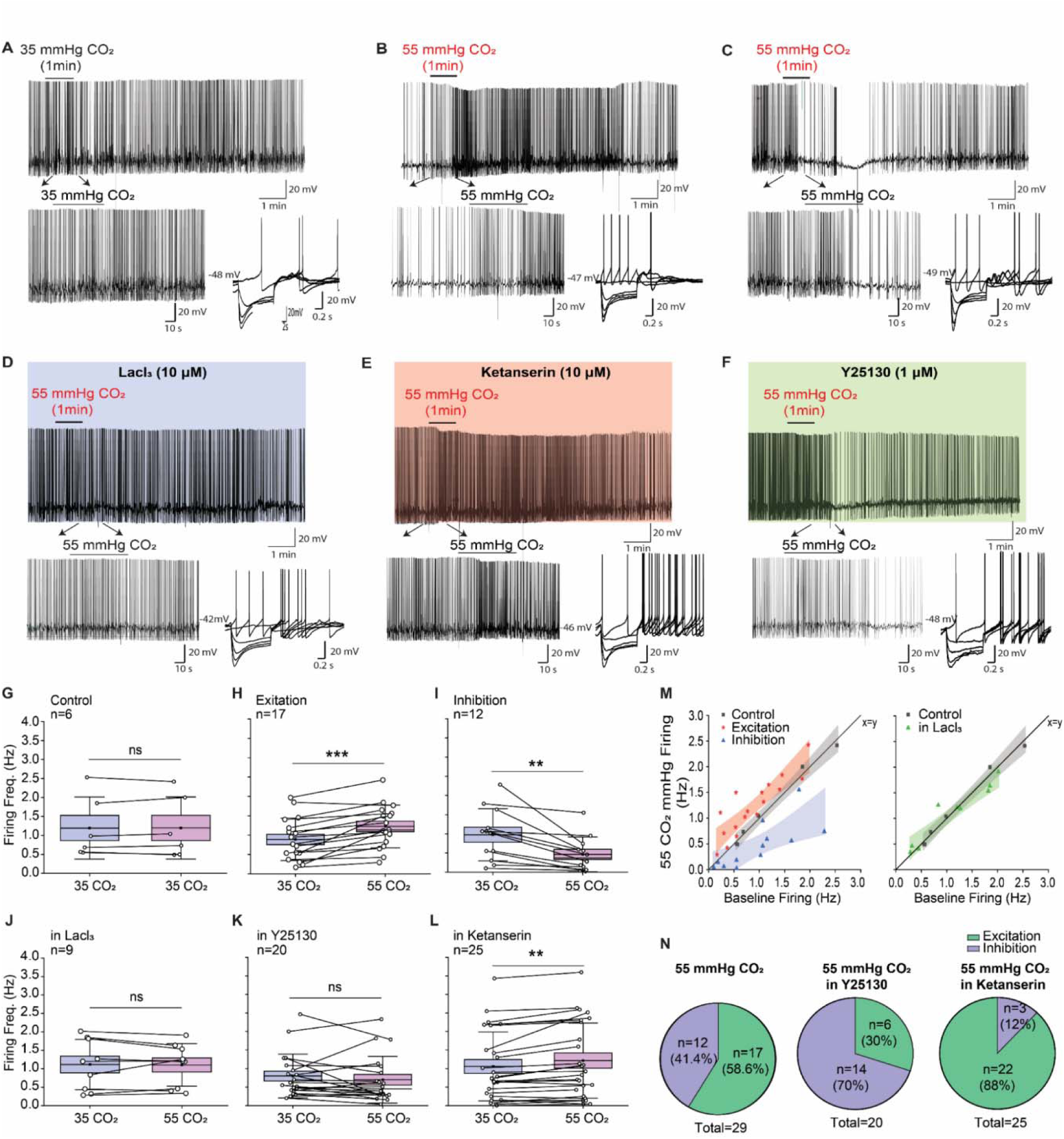
Brief PCO_2_ elevation alters VTA DA firing via 5-HT receptors. Whole-cell current-clamp recordings were obtained from VTA DA neurons in aCSF equilibrated with 35 mmHg CO_2_ (baseline), followed by exposure to either 55 mmHg CO_2_ for 1 min or continued perfusion with 35 mmHg CO_2_ (control). **(A-F)** Representative voltage traces showing spontaneous firing before, during, and after perfusion with 55 mmHg CO_2_ (or continuous perfusion with 35 mmHg CO_2_ in control experiments) under different experimental conditions (upper panels). Section from upper panel at an expanded time base to illustrate changes in the firing pattern (lower left panel) and current-voltage relationship of the recorded neurons showing characteristics of VTA DA neurons (lower right panel). **(G-L)** Summary plots showing the effects of elevated PCO_2_ on neuronal firing rate under different experimental conditions. (**G)** Firing rate remained stable during continuous perfusion with 35 mmHg CO_2_. (**H**) Exposure to 55 mmHg CO_2_ increased firing rate in 17 of 29 neurons (58.6%). (**I**) The remaining 12 of 29 neurons (41.4%) exhibited a decrease in firing rate during 55 mmHg CO_2_ exposure. (**J**) The effects of elevated CO_2_ on firing rate were abolished in the presence of the hemichannel inhibitor LaCl_3_. (**K**) The increase in firing rate induced by elevated CO_2_ was eliminated by the 5-HT_3_ receptor antagonist Y25130. **L**) In the presence of the non-selective 5-HT_3_ receptor antagonist ketanserin, elevated CO_2_ produced a significant increase in firing rate. (**M)** Scatter plot comparing baseline firing rates (35 mmHg CO_2_) with firing rates during elevated PCO_2_ (55 mmHg CO_2_), illustrating the segregation of neurons into CO_2_-excited and CO_2_-inhibited populations. (**N)** Pie charts summarizing the proportions of neurons exhibiting increased or decreased firing during exposure to 55 mmHg CO_2_ alone (58.6% increased, 41.4% decreased), in the presence of Y25130 (30% increased, 70% decreased), and in the presence of ketanserin (88% increased, 12% decreased). Data are mean ± SEM. Statistical significance was assessed using paired t-tests or Wilcoxon signed-rank tests, as appropriate. ***P < 0.0001, **P < 0.02; ns, not significant.

Additionally, we repeated the experiment using cell-attached recording to confirm that the observed effects were not due to the dialysis of DA neurons in whole-cell recording (**Fig. S5)**. These findings suggest the existence of functionally distinct DA neuron populations that respond differently to elevated CO_2_. To test whether the observed effects of elevated PCO_2_ required hemichannel opening, we used the hemichannel blocker LaCl_3_ (*42, 43*). Pre-treating the slices with LaCl_3_ (200 µM) abolished both the excitatory and inhibitory responses by raising PCO_2_ (**Fig. 4D, J, M Right)**. This finding is consistent with activation of the presynaptic Cx26-containing hemichannels identified in **Fig. 1**. Altogether, these results suggest that hemichannels are required for the CO_2_ -induced effects on DA neuron firing.

The observation that elevated PCO_2_ altered DA neuron firing without affecting glutamatergic transmission suggested that a non-glutamatergic neurotransmitter may mediate this effect. Given the serotonergic innervation of the VTA and the presence of Cx26-containing terminals, we investigated whether elevated PCO_2_ promotes 5-HT release and released 5-HT then activates serotonergic receptors expressed on DA neurons to modulate their activity. VTA DA neurons express multiple 5-HT receptor subtypes (*44-48*) and different receptor activation could potentially explain different responses observed following PCO_2_ elevation. Activation of 5-HT_3_ receptors is known to excite VTA DA neurons (*47-49*), encouraging us to test whether these receptors mediate the excitatory effects of elevated PCO_2_. In the presence of Y25130 (*38*), a selective 5-HT_3_ receptor antagonist, only 6 out of 20 cells (∼30% compared to ∼58.6% in 55 mmHg PCO_2_ alone) showed a modest increase in the firing frequency, compared with 17 of 29 neurons (∼59%) under elevated PCO_2_ alone (**Fig. 4 F, K, M**). The remaining 14 (∼70%) neurons showed a reduction in firing rate (**Fig. 4 F, K, M).** These findings indicate that the excitatory effects of elevated PCO_2_ on VTA DA neurons are largely mediated by activation of 5-HT_3_ receptors.

VTA DA neurons also express 5-HT_2A/C_ receptors, which produce an inhibitory effect (*50, 51*). To confirm whether these receptors regulate the inhibitory effects of elevated PCO_2_, recordings were repeated in the presence of the non-selective 5-HT_2A/C_ receptor antagonist ketanserin. In the presence of ketanserin, 22 of 25 neurons (∼88%, compared with ∼59% under elevated PCO_2_ alone) responded to 55 mmHg PCO_2_ with an increase in firing rate, whereas only 3 neurons (∼12%) exhibited a modest decrease in firing rate (**Fig. 4E, L, M**). Consequently, the overall effect of elevated PCO_2_ in the presence of ketanserin was a robust increase in DA neuron firing (**Fig. 4L)**. These findings indicate that the inhibitory effects of elevated PCO_2_ on VTA DA neurons are primarily mediated by activation of 5-HT_2A/C_ receptors. As subpopulations of VTA DA neurons might exhibit opposing responses, we compared their electrophysiological properties. No significant differences were detected between neurons that were excited or inhibited by elevated PCO_2_ (**Fig. S6A-I**). Taken together, these results show that a short, moderate CO_2_ challenge (55 mmHg for 1 min) alters the excitability of VTA DA neurons through activation of 5-HT_3_ and 5-HT_2A/C_ receptors.

To obtain independent evidence that elevated PCO_2_ promotes 5-HT release, we next examined its effects on hyperpolarization-activated cyclic nucleotide-gated (HCN) channels. VTA DA neurons express HCN channels that produce a prominent sag in response to hyperpolarising current steps (*52-55*). Because activation of 5-HT_2A/C_ receptors reduces HCN channel-mediated sag currents in VTA DA neurons, elevated PCO_2_ would be expected to inhibit the sag if it promotes 5-HT release. Increasing PCO_2_ (for 5 minutes) significantly inhibited sag amplitude and this inhibition was blocked by the hemichannel blocker LaCl_3_, by the 5-HT_2A/C_ antagonist ketanserin, by the selective 5-HT_2A_ receptor antagonist R96554 (**Fig. S7**). This inhibition could be partially reversed after prolonged wash (> 40 minutes) (**Fig. S8A-B**) and was mimicked by the direct application of 5-HT (**Fig. S8C)**. This further supports our hypothesis that CO_2_-evoked channel-mediated release of 5-HT can act on DA neurons to alter their excitability.

### Cx26 hemichannels are permeable to 5-HT

Our findings suggested that 5-HT may permeate Cx26 hemichannels opened by elevated PCO_2_. To test this directly, HeLa DH cells were co-transfected with Cx26 and the genetically encoded 5-HT sensor GRAB_5-HT_ (*56*). As HeLa cells do not make or store significant levels of 5-HT, the cells were first loaded with 5-HT by perfusing 5-HT (3 mM) into the cells in 55 mmHg PCO_2_ aCSF, a condition that opens Cx26 hemichannels. The PCO_2_ was then reduced to 20 mmHg to close hemichannels and trap 5-HT within the cells. Following washout of extracellular 5-HT, hemichannels were reopened by reapplying 55 mmHg PCO_2_ aCSF. If Cx26 hemichannels are permeable to 5-HT, reopening the channels should allow intracellular 5-HT to exit the cells and activate the GRAB_5-HT_ sensor, resulting in an increase in fluorescence (**Fig. 5A**). At the end of each experiment, the GRAB_5-HT_ sensor was calibrated by application of 10 µM 5-HT (**Fig. 5B**). Following the loading protocol, reopening Cx26 hemichannels with 55 mmHg PCO_2_ produced a robust increase in GRAB_5-HT_ fluorescence, indicating the release of intracellular 5-HT **(Fig. 5B and C**). This response was dependent on Cx26 expression, as cells transfected with GRAB_5-HT_ alone showed no increase in fluorescence upon PCO_2_ elevation (**Fig. 5C**). These results demonstrate that 5-HT can permeate open Cx26 hemichannels and support the feasibility of a mechanism in which elevated PCO_2_ opens Cx26 hemichannels on serotonergic terminals, thereby promoting 5-HT release.

**Fig. 5.**
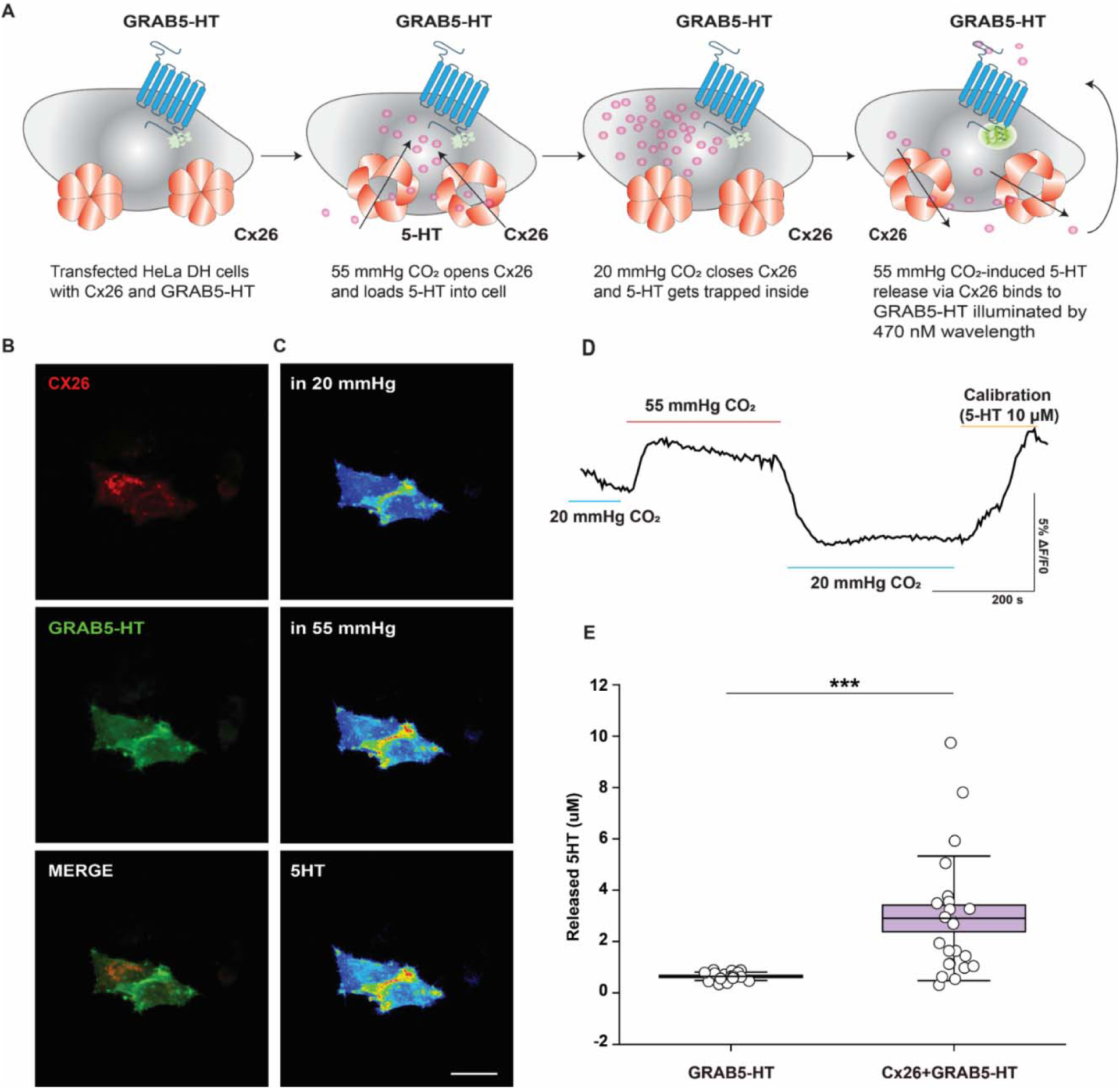
Opening of Cx26 hemichannels by raised PCO2 allows permeation of 5HT. **(A)** Schematic of the experimental protocol: Hela DH cells were co-transfected with Cx26 and the genetically encoded 5-HT sensor GRAB_5-HT_. Increasing PCO_2_ to 55 mmHg fully opens Cx26 hemichannels, allowing extracellular 5-HT to enter the cells. Hemichannels were then closed by reducing PCO_2_ to 20 mmHg, thereby trapping intracellular 5-HT. Following washout of residual extracellular 5-HT, PCO_2_ was increased again to 55 mmHg to reopen Cx26 hemichannels. If 5-HT is permeable through Cx26, its release from the cells is detected by GRAB_5-HT_ fluorescence during 470 nm excitation. (**B)** Representative images showing the expression of Cx26 (Red) and GRAB_5-HT_ (Green) in Hela DH cell. (**C-D)** Representative images from panel B showing GRAB_5-HT_ responses during exposure to 20 mmHg CO_2_ (blue bar), 55 mmHg CO_2_ (red bar), and 10 μM 5-HT (yellow bar). Scale bar: 20 μm. (**E)** Quantification of GRAB_5-HT_ fluorescence responses evoked by exposure to 55 mmHg CO_2_. Cells expressing GRAB_5-HT_ alone (n = 18 cells) were compared with cells co-expressing GRAB_5-HT_ and Cx26 (n = 22 cells from four independent transfections). Exposure to 55 mmHg CO_2_ produced a significantly larger increase in GRAB_5-HT_ signal in cells expressing both Cx26 and GRAB_5-HT_ than in cells expressing GRAB_5-HT_ alone, indicating Cx26-mediated release of 5-HT. Data are mean ± SEM. Statistical significance was assessed using a Mann–Whitney test. ***P < 0.0001.

### Hypercapnia activates VTA DA neurons

Our in vitro data suggest that an increase in PCO_2_ (hypercapnia) potentially activates VTA dopamine neurons in vivo. To test this, wild-type (WT) mice were exposed to either normoxic conditions or hypercapnia (9% CO_2_) for 30 min. Neuronal activation was then assessed by measuring expression of cFos, a widely used marker of neuronal activity (**Fig. 6A**). Both age groups were studied: juveniles (3–4 weeks old) and adults (3–4 months old). No significant differences were detected in the electrophysiological properties of VTA DA neurons between the two age groups (**Fig. S9A-I**), indicating that age is unlikely to contribute to any differences observed in vivo experiments.

**Fig. 6.**
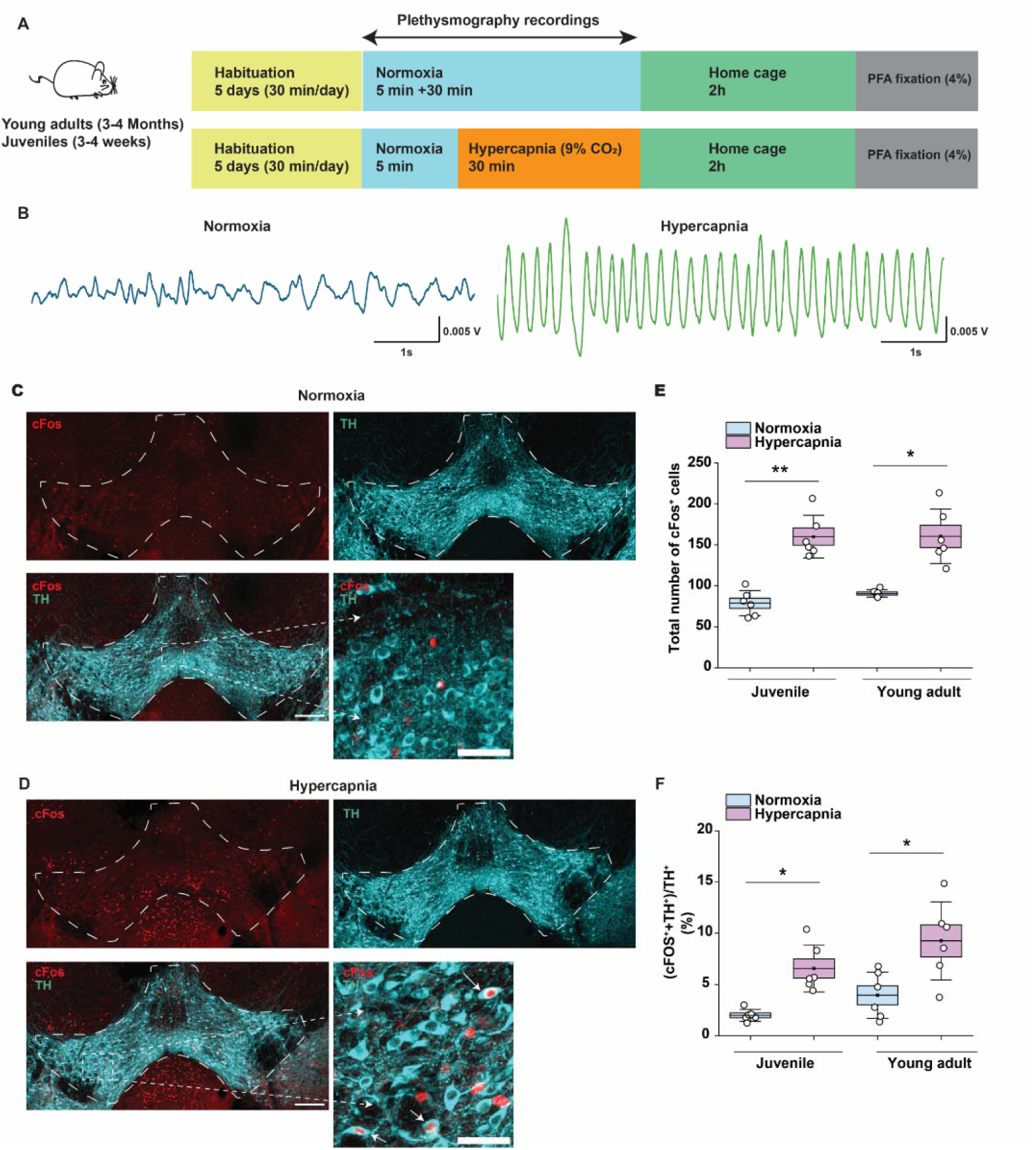
Hypercapnia activates DA neurons in the VTA. **(A)** the *in vivo* hypercapnic challenge protocol. (**B)** Representative breathing traces recorded during normoxia (slow breathing) and hypercapnia (faster, deeper breathing), demonstrating that hypercapnic exposure robustly alters respiratory patterns. (**C-D)** Representative images illustrating the cFos (red) expression and TH (cyan) expression in the VTA following normoxia **(C)** and hypercapnia **(D)**. White dashed outlines indicate the VTA with and white arrows highlight cFos^+^/TH^+^ colocalization (scale bars 200 µm; scale bar for cropped images 50 µm). (**E)** Quantification of cFos^+^ cells in the VTA of juvenile and adult mice exposed to normoxia or hypercapnia. Hypercapnic exposure significantly increased the total number of cFos^+^ cells in both age groups compared with normoxic controls. (**F)** Percentage of TH^+^ neurons expressing cFos (cFos^+^TH^+^/TH^+^). Hypercapnia significantly increased the proportion of activated DA neurons in both juvenile and adult mice relative to normoxia. Data are mean ± SEM. Statistical significance was assessed using two-way ANOVA followed by Tukey’s multiple-comparisons test. *P < 0.05, **P < 0.02.

Following exposure to hypercapnia, breathing frequency and tidal volume increased, confirming a robust physiological response to elevated CO_2_ (**Fig. 6B).** We then checked neuronal activation by counting the total number of cFos-positive cells (cFos^+^) within the VTA. Hypercapnia significantly increased the number of VTA cFos^+^ cells in both juvenile and adult mice compared with their normoxic controls (**Fig. 6C-E**). We then quantified the proportion of TH^+^ neurons that co-express cFos (cFos^+^TH^+^) within the VTA. We found that hypercapnia significantly increased the proportion of cFos^+^TH^+^ neurons in both juvenile and adult mice **(Fig. 6F)**, demonstrating that hypercapnia activates VTA DA neurons.

### Loss of Cx26 CO**_2_** sensing from DR neurons delays arousal

The presence of Cx26 in the DR (**Fig. 2D-E**), together with the increased firing rate of DR neurons during exposure to 55 mmHg CO_2_ for 1 min (**Fig. S10**), suggests that DR neurons are directly responsive to CO_2_. Because DR 5-HT neurons project to the VTA, it is plausible that CO_2_ may modulate VTA DA activity directly through the Cx26-expressing DR neurons. To test this hypothesis, we engineered a dominant negative Cx26 virus under the TPH2 promoter, ensuring expression in DR 5-HT neurons. This manipulation abolished CO_2_ sensitivity of Cx26 in these neurons. After injecting either rAAV-TPH2-Cx26WT-STOP-IRES-Clove-rWPRES (Cx26^WT^) or rAAV-TPH2-Cx26DN-STOP-IRES-Clover-WPRES (Cx26^DN^) into the DR of C57BL/6 mice (**Fig. 7A and Fig. S11**), we found that TPH2-promoted virus targeted the majority of quantified 5-HT neurons in the DR (Cx26^WT^: ∼ 62% and Cx26^DN^: ∼ 65%) (**Fig. 7B** and **Fig. S12**). Following 5 days of habituation, mice were exposed to a 30 s puff of 6% CO_2_ while EEG activity was recorded (**Fig. 7C**). Hypercapnic arousal was assessed by measuring arousal latency, defined as the time required to transition from sleep to wakefulness following CO_2_ exposure (**Fig. 7D**). In the Cx26^WT^ -expressing mice, arousal latency remained stable across weeks 3, 4, and 5 (**Fig. 7E**), indicating that repeated brief CO_2_ exposures do not necessarily desensitize the hypercapnic arousal reflex. Additionally, there is no significant difference between control (non-virus injection) and Cx26^WT^, demonstrating that Cx26^WT^ virus itself does not change arousal latency (**Fig. 7E**).

**Fig. 7.**
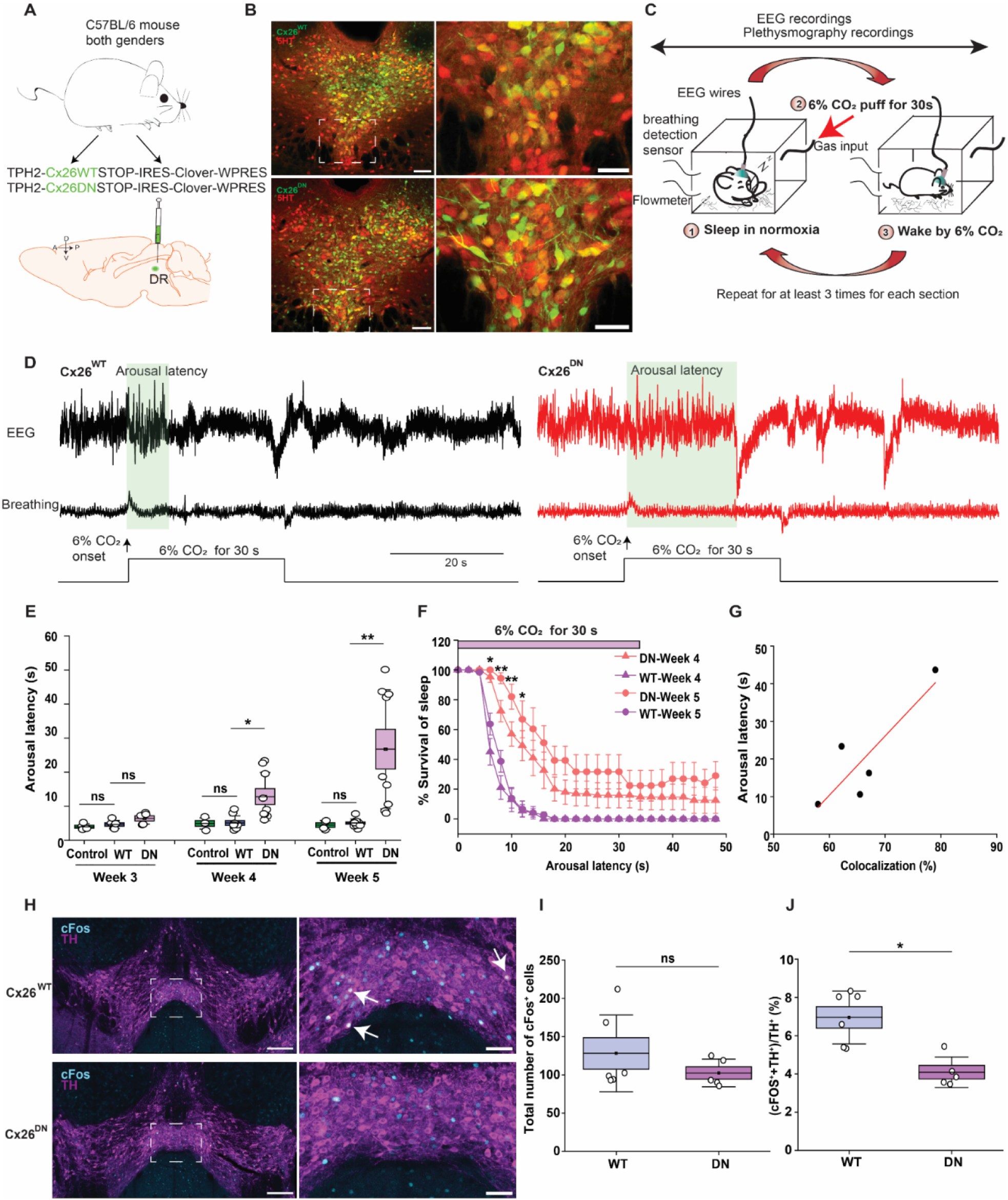
Loss of Cx26 CO_2_ sensing from DR neurons delays hypercapnic arousal. **(A)** Schematic illustrating injection of TPH2-promoted Cx26^WT^ and Cx26^DN^ virus injection into mouse DR. (**B)** Representative images showing the expression of Cx26^WT^ and Cx26^DN^ in 5-HT-labelled serotonergic neurons of the DR. (**C)** Experimental paradigm for CO_2_ exposure and EEG recording. (**D)** Representative EEG and respiratory traces recorded during exposure to 6% CO_2_ in Cx26^WT^ and Cx26^DN^ expressing mice. (**E)** Quantification of arousal latency in response to 6% CO_2_ at weeks 3, 4, and 5 after viral injection. No significant differences were observed among groups at week 3. In contrast, mice expressing Cx26^DN^ showed significantly longer arousal latency compared with Cx26^WT^ group at weeks 4 and 5. (**F)** Sleep survival plot showing the proportion of sleeping animals following 6% CO_2_ exposure at weeks 4 and 5. Significant differences between Cx26WT and Cx26DN groups were observed at 6, 8, 10, and 12 s after CO_2_ onset. **(G)** Correlation between the proportion of virus (Green)-transduced serotonergic (5HT^+^, red) neurons and arousal latency following CO_2_ puffs (*n* = 5 animals), linear regression analysis showed an R-Square value of 0.752. (**H)** Representative images showing cFos expression (cyan) in VTA DA neurons (TH, magenta) following 30 minutes exposure to 9 % CO_2_ in mice expressing Cx26^WT^ or Cx26^DN^ in DR. (**I)** Quantification of cFos^+^ cells in the VTA. Expression of Cx26^DN^ significantly reduced the proportion of activated DA neurons compared with Cx26^WT^ controls (Cx26^WT^, n = 6 animals; Cx26^DN^, n = 5 animals) (**J)** Quantification of the percentage of TH^+^ neurons expressing cFos^+^ in the VTA. Expression of Cx26^DN^ significantly reduced the proportion of activated DA neurons compared with Cx26^WT^ controls (Cx26^WT^, n = 6 animals; Cx26^DN^, n = 5 animals). Data are presented as mean ± SEM. Statistical significance was assessed using two-way ANOVA followed by Tukey’s multiple-comparisons test for panel F and a Mann–Whitney test for panel I. *P < 0.05, **P < 0.002.

In contrast, mice expressing Cx26^DN^ showed a progressive increase in arousal latency beginning at week 4 (**Fig. 7E**), suggesting that prolonged expression of Cx26^DN^ is required before its functional effects become apparent. The significant increase in arousal latency at weeks 4 and 5 in Cx26^DN^-expressing mice indicates that Cx26-mediated CO_2_ sensing in DR 5-HT neurons contributes importantly to the hypercapnic arousal reflex (**Fig. 7E**). Furthermore, sleep survival analysis revealed a slower decline in the proportion of sleeping mice in the Cx26^DN^-expressing mice compared with Cx26^WT^ controls, indicating delayed arousal latency following CO_2_ exposure (**Fig. 7F**). The severity of the arousal latency impairment was positively correlated with the proportion of transduced 5-HT neurons in Cx26^DN^ expressing mice (**Fig. 7G**).

To address whether this delayed arousal response alters downstream activity in VTA DA neurons, we exposed these animals to 9% CO_2_ for 30 minutes and quantified cFos expression. We counted both the total number of cFos^+^ cells and the proportion of activated DA neurons (the number of cFos^+^TH^+^ cells out of the total TH^+^ cells) in the VTA. Although the total number of cFos-positive cells was unchanged between groups (**Fig. 7H, I)**, Expression of Cx26^DN^ reduced the number of cFos^+^TH^+^ neurons compared with Cx26^WT^ mice (**Fig. 7H, J)**. These findings indicate that Cx26-dependent CO_2_ sensing in DR serotonergic neurons contributes to the activation of VTA DA neurons during hypercapnia and plays an important role in promoting arousal in response to elevated CO_2_.

## DISCUSSION

Our results demonstrate that Cx26 hemichannels are expressed in both the soma and VTA projecting axon terminals of DR serotonergic neurons. Raised levels of PCO_2_ open Cx26 hemichannels present in these terminals, allowing the presynaptic release of 5-HT. The released 5-HT subsequently activates two subtypes of 5-HT receptor (5-HT_3_ and 5-HT_2A/C_) expressed on VTA DA neurons to either increase or decrease their firing rate. Removal of Cx26 sensitivity to CO_2_ from DR 5-HT neurons, by using a dominant negative Cx26 viral construct, significantly delayed arousal in response to CO_2_ puffs and decreased activity of VTA DA neurons.

### Effects of raised PCO_2_ on neurotransmission in the VTA

The opening of Cx26 hemichannels, by increased PCO_2_, expressed in SERT-positive nerve terminals could potentially contribute to changes in exocytotic neurotransmission (of glutamate and/or 5-HT). This is because Cx26 hemichannels are permeable to Ca^2+^and their opening would likely elevate the levels of Ca^2+^ within the terminals (*57, 58*). However, we found that increases in CO_2_ had no effect on the frequency or amplitude of either miniature postsynaptic currents (PSCs) or isolated miniature AMPA receptor-mediated miniature EPSCs. As Ca^2+^ dependent exocytosis is triggered by Ca^2+^ influx via voltage gated Ca^2+^ channels (VGCCs) that are concentrated at active zones (*59*), we speculate that the hemichannels could be located relatively distant from the active zone or alternatively any Ca^2+^ influx via the hemichannels is quickly removed by intracellular Ca^2+^ buffers prior to forming the microdomains required for exocytosis (*60*). Although there was no change in mPSC frequency, our electrophysiological recordings showed that increases in PCO_2_ produced the activation of two subtypes of 5-HT receptors that modulate DA neuron excitability. As there is no evidence so far showing the presence of serotonergic neurons within the VTA, it is plausible that CO_2_-induced 5-HT release most likely originated from presynaptic terminals projecting from the DR. While this presynaptic release could be via exocytosis of 5-HT-containing synaptic vesicles, we suggest that a large component is via direct channel-mediated release for the following reasons: Firstly, Cx26 is expressed in the serotonergic terminals of the VTA, and in their projecting neurons in the DR; Secondly, we have shown that 5-HT can freely permeate through Cx26 hemichannels opened by raised PCO_2_. Thirdly, the CO_2_-evoked release of 5-HT, that modulates the excitability of VTA dopaminergic neurons in the VTA, could be blocked by a hemichannel blocker LaCl_3_.

### Channel-mediated transmitter release

Channel-mediated volume transmission via ATP from glial cells has been described in several contexts such as the developing retina and cortex, chemosensory regulation of breathing, chemosensory transduction in the hypothalamus (*61*). However, a neuronal form of channel mediated transmission has only been described in taste buds and in the airways. In the taste pathway, CALHM1/3 channels open in response to transmembrane depolarisation allowing the release of ATP to diffuse a short distance (∼30 nm) to activate postsynaptic receptors. The very close proximity of the source of ATP (an “atypical” mitochondrion), the release channels (CALHM1/3), and the postsynaptic P2X2/3 receptors has led to the term “channel synapse” (*62-64*). Recently, CALHM1/3 channel synapses were also found in throat chemosensory cells that participate in airway protective programs (*65*). It is currently unclear whether such channel synapses occur more widely in the brain. While this remains an open question, our findings in the VTA suggest a new but related concept: the “channel co-synapse”. Our data suggests the existence of Cx26 in presynaptic terminals, that open to allow nerve terminal autonomous release of 5-HT in response to CO_2_ independently of activation of the neuronal cell body. This channel-mediated release would occur in parallel with the conventional exocytic release of 5-HT presumably controlled by action potential firing that is initiated in the DR cell bodies. We currently lack detailed morphological evidence to know whether the intricate juxtaposition of the essential components of transmission described for channel synapses in the taste cells also occurs in the serotonergic terminals of the VTA. However, it is possible that channel co-synapses could be widespread. Because these co-synapses could occur in parallel with conventional synaptic transmission, they will only be discovered by specific experiments that directly test this hypothesis.

### CO_2_-dependent 5HT release has dual effects on VTA DA neurons

Previous studies showed that VTA DA neurons express both 5-HT_3_ (*38, 47, 48*) and 5-HT_2A/C_ receptors (*44-46, 66, 67*). In our work, we found that a brief (1 minute) CO_2_ challenge could result in either inhibition or excitation of DA neurons. Application of receptor antagonists revealed that 5-HT_3_ receptors produced the excitatory effects (increasing firing) of CO_2_, whereas 5-HT_2A/C_ receptors mediated the inhibitory effects (reducing firing) of CO_2_. This suggests that the overall effect of raised PCO_2_ on individual DA neurons will depend on the relative degree of activation of 5-HT_3_ and 5-HT_2A/C_ receptors. This may depend on the amount of 5-HT released, as low concentrations of 5HT will preferentially bind to 5-HT_2A/C_ due to their higher affinity compared to 5-HT_3_ receptors (*68*). To our current knowledge, the expression pattern and distribution of 5-HT receptors on VTA DA neurons remains unclear, we speculate that the proximity to the 5HT release sites determines which receptors are preferentially activated.

### Limitations of the study

Our findings suggest that Cx26 hemichannels contribute to CO_2_ dependent signalling in both the VTA and dorsal raphe. However, we did not fully define the structural organization or molecular composition of the proposed Cx26 hemichannel co synapses. High-resolution anatomical approaches will be required to confirm the precise arrangement of Cx26 relative to serotonergic terminals and 5-HT receptors on dopaminergic postsynaptic sites. In addition, while we demonstrate CO_2_ mediated modulation of hypercapnic arousal in the dorsal raphe, our experiments do not yet establish the full range of functional consequences across the diverse Cx26 expressing-5 HT subpopulations within this nucleus and their long-range projections. Future work will need to characterize subtype or division expression of Cx26 in DR, their projection targets and behavioural outcomes to fully understand its contribution to hypercapnic arousal.

## MATERIALS AND METHODS

### Study design

The objective of this study was to determine whether Cx26 hemichannels expressed by dorsal raphe serotonergic neurons contribute to hypercapnic arousal. We combined immunostaining, electrophysiological, imaging, viral-genetic, and behavioral approaches to identify CO_2_ -sensitive Cx26 hemichannels, examine their role in serotonin release and ventral tegmental area dopamine neuron activity, and assess their contribution to arousal responses during hypercapnia. Data analysis was performed blind whenever possible.

### Animals

All animal procedures were approved by the Animal Welfare and Ethical Review Board (AWERB) of the University of Warwick and carried out in accordance with the Animals (1986) Scientific Procedures Act of the UK under the authority of Licence PP7458325. For electrophysiological recordings, C57BL/6J mice (both sexes) between 3-5 weeks and, 9–10 weeks-old were used.

### Acute brain slice preparation

In accordance with the Animals (1986) Scientific Procedures Act of the UK, 3-4 week old (both sex) mice were killed by cervical dislocation and decapitated. The brain was quickly removed and sagittal brain slices (350 µm) were cut using a vibratome (Microm HM 650V microslicer) with ice-cold cutting solution composed of (in mM: 85 NaCl, 2.5 KCl, 0.5 CaCl_2_, 1.25 NaH_2_PO_4_, 24 NaHCO_3_, 25 glucose, and 75 sucrose; pH was adjusted to 7.4, 290–300 mOsm) bubbled with 95% O_2_ and 5% CO_2_. The slices were then stored in aCSF composed of (in mM: 127 NaCl, 1.9 KCl, 2 MgCl_2_, 2 CaCl_2_, 1.2 KH_2_PO_4_, 26 NaHCO_3_, 10 D-glucose, pH 7.4, 290–300 mOsm when bubbled with 95% O_2_ and 5% CO_2_) maintained at 33 °C for 1-6 hours.

For slice preparation from 9-10 week old mice, mice were deeply anaesthetized with isoflurane, and perfused transcardially with 50 ml of ice cold NMDG aCSF (in mM: 93 NMDG, 2.5 KCl, 1.2 aH_2_PO_4_, 30 NaHCO_3_, 20 HEPES, 25 glucose, 5 Na-ascorbate, 2 thiourea, 3 Na-pyruvate, 12 N-acetyl-L-cysteine, 10 MgSO_4_, 0.5 CaCl_2_, pH adjusted to 7.3-7.4 with 10 M HCl). Mice were rapidly decapitated, and heads were submerged in NMDG-aCSF. Sagittal slices (200 μm) were sliced with vibratome (Microm HM 650V microslicer) in ice-cold NMDG-aCSF. The slices were incubated in aCSF composed of (in mM: 127 NaCl, 1.9 KCl, 2 MgCl_2_, 2 CaCl_2_, 1.2 KH_2_PO_4_, 26 NaHCO_3_, 10 D-glucose, pH 7.4, 290–300 mOsm bubbled with 95% O_2_ and 5% CO_2_) at 33 °C.

### Whole-cell patch-clamp recording

A slice was transferred to the recording chamber and perfused with oxygenated aCSF at a rate of 2.5-3 ml/min at 33 ± 0.5°C. Neurons in the VTA were visualized using differential interference contrast (DIC) optics with an Olympus BX151W microscope (Scientifica, Bedford UK). Cell attached and whole-cell patch clamp recordings were made using patch pipettes (4-7 MΩ) manufactured from thick-walled borosilicate glass (Harvard Apparatus, Edenbridge UK) containing (mM): 135 potassium gluconate, 7 NaCl, 10 HEPES, 0.5 EGTA, 10 phosphocreatine, 2 MgATP, 0.3 NaGTP, 293∼297 mOSM, pH 7.2). Some of the neurons were filled with 3 mM biocytin via the patch pipette for immunohistochemistry. Current and voltage clamp recordings were obtained using an Axon Multiclamp 700B amplifier (Molecular Devices, USA) and digitised at 20 KHz. Data acquisition and analysis was performed using pClamp10 software (Molecular Devices, USA). The bridge balance was monitored throughout the experiments and any recordings where it changed by >20% were discarded. The electrophysiological parameters of VTA dopamine neurons were extracted using step current-voltage relationships with 50 pA steps of 1s duration, starting from -200 pA and going up to +50 pA.

### Measuring spontaneous synaptic transmission

Spontaneous postsynaptic currents were recorded at a holding potential of -70 mV, filtered at 3 KHz and digitised at 20 KHz. To record miniature post synaptic currents (mPSCs), tetrodotoxin (TTX, 1 mM) was present in the aCSF solutions to block action potentials. To isolate AMPA receptor-mediated miniature excitatory post synaptic currents, the GABA_A_ receptor antagonist picrotoxin (PTX: 50 mM) and the NMDA receptor antagonist L689,450 (5 mM) were also present. For analysis of the postsynaptic currents, the ClampFit template search method was used (pClamp, Molecular Devices). A template was constructed for at least 100 events and a template match stringency of 4 was used, with events manually accepted following visual inspection.

### Isohydric solutions

Three different aCSF solutions were used. For physiological conditions, 35 mmHg CO_2_ aCSF was used, composed of (in mM): 124 NaCl, 26 NaHCO_3_, 1.25 NaH_2_PO_4_, 3 KCl, 10 D glucose, 1 MgSO_4_, 2 CaCl_2_, bubbled with 95% O_2_ and 5% CO_2_ at a final pH of ∼7.4. To mimic hypercapnia, 55 mmHg CO_2_ aCSF was used composed of (in mM): 100 NaCl, 50 NaHCO_3_, 1.25 NaH_2_PO_4_, 3 KCl, 10 D glucose, 1 MgSO_4_, 2 CaCl_2_. To mimic hypocapnia, 20 mm Hg CO_2_ aCSF was used composed of in (mM: 140 NaCl, 10 NaHCO_3_, 1.25 NaH_2_PO_4_, 3 KCl, 10 D glucose, 1 MgSO_4_, 2 CaCl_2_). The pH of both hypercapnic and hypocapnic solutions was adjusted to match physiological 35 mmHg CO_2_ aCSF by balancing CO_2_ and O_2_ (*25, 29, 34*). Thus, any changes in PCO_2_ do not change extracellular pH, which remained constant.

### Transcardial perfusion for FITC-dye loading

For FITC loading experiments, mice aged between 2-3 months (both sexes) were used. The FITC (50 mM) was dissolved into both hypercapnic (55 mmHg CO_2_) and hypocapnic (20 mmHg CO_2_) solutions. The animals were first culled with an overdose of isoflurane and then transcardial perfusion was performed. The perfusion started with 10 ml of 55 mm Hg CO_2_ with FITC at a rate of 2.5 ml/min, followed by 10 ml of 20 mmHg CO_2_ with FITC. For the control group, 20 ml of 20 mmHg CO_2_ with FITC was perfused. At the end, 10 ml of 4% PFA was used for post fixation overnight at 4°C. The brains were sliced into 50 mm thick slices using a vibratome (Microm HM 650V microslicer) in PBS for immunofluorescence labelling.

### Immunohistochemistry

Mice aged between 2-3 months (both sexes) were culled with an overdose of isoflurane and then transcardially perfused with 0.9% saline and 4% PFA. After post-fixation with 4% PFA, the brains were sliced into 50 mm thick slices using a vibratome (Microm HM 650V microslicer) in PBS. The free-floating slices (with or without FITC) in PBS were washed 3 times by 1x PBS at room temperature (RT) on the shaker. The slices were then incubated in blocking solution (4% BSA, 0.1% Triton x-100, and 0.01% sodium azide in PBS) for 2 hr. Subsequentially, the slices were incubated with antibodies at 4°C for 48 hr. The primary antibodies: Chicken anti-Tyrosine hydroxylase (1:1000), Sheep anti-Tyrosine hydroxylase (1:1000), Mouse anti-Aquaporin 5 antibody (1:200), Rabbit anti-Connexin 26 (1:500), Mouse anti-Connexin 26 (1:250), Guinea pig anti-Sert (1:3000), Guinea pig anti-Vglut2 (1:250); Mouse anti-Vglut3 (1:100), Rabit anti-c-fos (1:1000). After washing steps, the slices were incubated with secondary antibodies diluted in the blocking solution (composition outlined above) for 3-4 hr at room temperature. Secondaries: Donkey anti-Chicken IgY (H+L) Highly Cross Adsorbed Secondary Antibody, Alexa Fluor™ 647 (1:250), Goat anti-Chicken IgY (H+L) Secondary Antibody, Alexa Fluor™ 488 (1:250), Donkey anti-Sheep IgG (H+L) Cross-Adsorbed Secondary Antibody, Alexa Fluor™ 594 (1:250), Goat anti-Rabbit IgG (H+L) Cross-Adsorbed Secondary Antibody, Alexa Fluor™ 488 (1:250), Donkey anti-Mouse IgG (H+L) Highly Cross-Adsorbed Secondary Antibody, Alexa Fluor™ 488 (1:250), Goat anti-Mouse IgG (H+L) Cross-Adsorbed Secondary Antibody, Alexa Fluor™ 594 (1:250), Donkey anti-Mouse IgG (H+L) Highly Cross-Adsorbed Secondary Antibody, Alexa Fluor™ 647 (1:250), Donkey anti-Rabbit IgG (H+L) Highly Cross-Adsorbed Secondary Antibody, Alexa Fluor™ 594 (1:250), Donkey anti-Rabbit IgG (H+L) Highly Cross-Adsorbed Secondary Antibody, Alexa Fluor™ 488 (1:250), Goat anti-Guinea Pig IgG (H+L) Highly Cross-Adsorbed Secondary Antibody, Alexa Fluor™ 647 (1:250). After incubation, the samples were quickly washed again and mounted on the Super Frost Plus microscope slides (25 x 75 x 1 mm) for confocal microscope (Zeiss LSM880) visualization. The Z stack images were obtained using a 63x objective with 2.5 zoom (Zeiss LSM880) for the quantification of FITC-loaded puncta. To quantify the puncta, masks were generated based on threshold and used to measure the number, mean intensity, and volume of the FITC puncta on the binary mask generated in ImageJ. Both analysis and experiments were performed blind. For cFos quantification, Z-stack plus tile images were taken with a 25x objective (Zeiss LSM880). To quantify cFos positive cells, the same intensity was applied to the images from normoxia and hypercapnia groups performed on the same day. Python cellpose together with ImageJ were used to count the total number of cFos and the number of cFos^+^TH^+^ cells in the VTA.

### RNAscope in situ hybridization

After post-fixation, brains were cryoprotected in 30% sucrose containing PBS two nights. Brains were mounted with OCT Tissue-Tek and sliced with cryostat. Coronal sections (15 mm) were collected on SuperFrost Plus slides. The RNAcope method was performed as previously described with a modification (*69*). Sections were kept at RT overnight and then incubated at 40°C for 5 min to dry. To dehydrate the sections, a sequence of ethanol serial dilutions from 50%, 75%, 95%, and 100% (5 min for each dilution) were performed. The dehydrated sections were further incubated in hydrogen peroxide (H_2_O_2_) at RT for 10 min, followed by briefly washing in distilled water (dH_2_O). Hydrophobic barrier was drawn by a Pap Pen (Vector Labs). Subsequentially, sections were treated with protease III (ACD, #322336) at RT for 20 min. Following two times rinses in dH_2_O (2 min each time), sections were incubated in GJB2 probe (#518881-C2, ACD Bio) and SLC17A6 (#319171-C3, ACD Bio) with a dilution of 1:50, SLC32A1 (#319191-C1, ACD Bio) with full concentration at 40°C in the HybEZTM oven (ACD) for 2h. After brief wash in 1x RNAscope washing buffer (ACD, #310091), a series of signal amplification steps were performed at 40°C in the HybEZTM, including Amp 1-FL-V2 (30 min), Amp 2-FL-V2 (30 min), and Amp 3-FL-V2 (15 min). Between amplification steps, 2 times washing (2 min each time) in 1x wash buffer was performed. Sections were further incubated in FL v2 HRP-C2 at 40°C for 15 min, followed by labelling with fluorophore-conjugated Tyramine plus (TSAP)-650 (1:1500) and 570 (1:1500) at 40°C for 30 min. Following 2 times wash in washing buffer (2 min each time), sections were incubated in HRP blocker at 40°C for 15 min and stained with DAPI using the reagent provided by the Fluorescent Multiplex Kit. Finally, sections were mounted by Prolong gold mounting medium and cover slipped for visualization. For the visualization, confocal microscope (Zeiss LSM880) was used.

### 5HT release experiments

Parental HeLa DH cells (ECACC 96112022) were cultured with low-glucose DMEM supplemented with 10% foetal bovine serum (FBS) and 5% penicillin/streptomycin. Cells were seeded onto coverslips at a density of 4x10^4^ cells per well of a 6-well plate. Cells were transiently transfected to co-express Cx26-pCAG-GS-mCherry and the genetically encoded 5HT sensor: pDisplay-GRAB_HTR6-1.0 was a gift from Yulong Li (Addgene plasmid # 180005; http://n2t.net/addgene:180005; RRID: Addgene180005) 48 hours prior to imaging. Cells were perfused with control aCSF until a stable baseline was reached, before perfusion with hypercapnic aCSF. Once a stable baseline was reached after solution change, cells were again perfused with control aCSF and when a stable baseline reached, recordings were calibrated by application of 5HT. All cells were imaged by epifluorescence (Scientifica Slice Scope, Cairn Research OptoLED illumination, 60x water Olympus immersion objective, NA 1.0, Hamamatsu ImagEM EM-SSC camera, Metafluor software). cpGFP in the sensor was excited by a 470 nm LED, with emission captured between 504-543 nm. Cx26 has a C-terminal mCherry tag, which is excited by a 535 nm LED and emission captured between 570-640 nm. Only cells expressing both cpGFP and mCherry were selected for recording, with cpGFP images acquired every 4 seconds. At least 3 independent transfections were performed with at least 2 coverslips per transfection. Analysis of all experiments was carried out in ImageJ. Images were opened as a stack and stabilised. ROIs were drawn around cells co-expressing both sensor and connexin. Median pixel intensity was plotted as normalised fluorescence change (ΔF/F_0_) over time to give traces of fluorescence change. Amount of analyte release was quantified as concentration by normalising to the ΔF/F_0_ caused by application of 10 µM 5HT.

### Viral injection and expression

rAAV-TPH2-Cx26WTSTOP-IRES-Clover-WPRES and rAAV-TPH2-Cx26DNSTOP-IRES-Clover-WPRES were customized by BrainVTA (Wuhan, China). The viruses were aliquoted and stored at −80°C before use. The surgery process was descried previously (*70*). 9–10 week-old C57BL6 animals both genders were anesthetized with isoflurane (4%; Piramal Healthcare Ltd., Mumbai, India) in pure oxygen (4 L·min−1). Anaesthesia was maintained with 0.5–2% isoflurane in pure oxygen (1 L·min−1) throughout the surgery. Mice received a pre-surgical subcutaneous injection of atropine (120 μg·kg−1; Westward Pharmaceutical Co., Eatontown, NJ, USA) and meloxicam (2 mg·kg−1; Norbrook Inc., Lenexa, KS, USA). Mice were placed in a digital stereotaxic frame (Kopf Instruments, Tujunga, CA, USA) on a heating pad (TCAT 2-LV: Physitemp, Clifton, NJ, USA), and body temperature was monitored at a minimum value of 33°C via a thermocouple. After removal of the skin, a small unilateral craniotomy was made above the DR (from bregma: AP: -4.5 mm; ML: 0 mm; DV: -2.8 mm). 100-150 nl virus was injected and the capillary was kept in place for 5 min before retraction.

### EEG implantation and recordings

After virus delivery, EEG electrodes were implanted as previously described (*70*). Briefly, four holes were drilled for two EEG recording electrodes (from Bregma AP: +1.0 mm, ML: ±1.5 mm), a ground and a reference electrode (from Bregma AP: -1.5 mm, ML: ±2.5 mm). EEG electrodes were made of PFA-coated silver wire with 0.254 mm diameter (Bilaney Consultants Ltd., UK). Silver wires were implanted and secured in position with SuperBond™ (Prestige Dental, Bradford, UK). Mice received subcutaneous injections of buprenorphine (100 μg·kg^−1^; Reckitt Benckiser, Slough, UK). After recovery from aesthesia, the mice were returned to their home cages. Mice were habituated in a custom-built plethysmograph chamber (*29*) for 5 days with EEG connections prior to sleep recordings. Sleep recordings were performed as previously described with modifications (*70*). The EEG signal, CO_2_ level and breathing responses were amplified and filtered using the NeuroLog system connected to a 1401 interface acquired on a computer using the Spike2 software with a built-in camera (Cambridge Electronic Design). EEG signals were sampled at 6250 Hz. Gas composition (O_2_ and CO_2_) was monitored by a gas analyser (Hitech Instruments, GIR250 Dual Sensor Gas Analyser). A 6% CO_2_ puff was delivered for 30 s once the mouse showed at least 20s of sleep in the chamber, defined by show wave EEG activity. Each animal received a minimum of two CO_2_ puffs per session, with at least 5 min interval between puffs. Recordings were done at 3, 4 and 5 weeks following viral injection. As the onset of CO_2_ delivery produced a characteristic pressure artefact in the breathing trace, the peak of this artefact was used to mark the precise time of CO_2_ onset across all of the recordings. Corresponding EEG traces were then aligned to this point to calculate arousal latency, defined as the transition from slow to fast EEG activity. The EEG data were independently analysed.

### Statistical analysis

Statistical tests were run in Microcal Origin 2024b. The number (n) of cells or mice for each experimental conditions, details of comparison, statistical methods (a normality test was performed to determine whether to use parametric methods for each graph) can be found in the relevant figure legends. All data are shown as mean ± SEM.

## Author contributions

Conceptualization: **L.H, N.D and M.J.W**

Methodology: **L.H, N.D and M.J.W**

Investigation: **L.H, J.B, A.L, J.B, J.P, E.L.H**

Analysis**: J.B**

Supervision: **N.D and M.J.W**

Writing—original draft: **L.H, N.D and M.J.W**

Writing—review & editing: **L.H, N.D and M.J.W**

## Data, code, and materials availability

All data supporting the findings of this study are available within the paper and its Supplementary Materials. The complete maps and annotated sequences of the recombinant AAV constructs used in this study are provided as Auxiliary Files for peer review and will be made publicly available upon publication. Additional data and materials are available from the corresponding author upon reasonable request.

## REFERENCES

1. B. W. Locke, J. P. Brown, K. M. Sundar, The Role of Obstructive Sleep Apnea in Hypercapnic Respiratory Failure Identified in Critical Care, Inpatient, and Outpatient Settings. Sleep Med Clin 19, 339–356 (2024).

2. C. Dave, S. Wharton, R. Mukherjee, B. M. Faqihi, R. A. Stockley, A. M. Turner, Development and Relevance of Hypercapnia in COPD. Can Respir J 2021, 6623093 (2021).

3. D. J. Eckert, M. K. Younes, Arousal from sleep: implications for obstructive sleep apnea pathogenesis and treatment. J Appl Physiol (1985) 116, 302–313 (2014).

4. J. E. Remmers, W. J. deGroot, E. K. Sauerland, A. M. Anch, Pathogenesis of upper airway occlusion during sleep. J Appl Physiol Respir Environ Exerc Physiol 44, 931–938 (1978).

5. S. Kaur, C. B. Saper, Neural Circuitry Underlying Waking Up to Hypercapnia. Front Neurosci 13, 401 (2019).

6. Y. Arima, S. Yokota, M. Fujitani, Lateral parabrachial neurons innervate orexin neurons projecting to brainstem arousal areas in the rat. Sci Rep 9, 2830 (2019).

7. S. Kaur, N. P. Pedersen, S. Yokota, E. E. Hur, P. M. Fuller, M. Lazarus, N. L. Chamberlin, C. B. Saper, Glutamatergic signaling from the parabrachial nucleus plays a critical role in hypercapnic arousal. J Neurosci 33, 7627–7640 (2013).

8. S. Kaur, R. De Luca, M. A. Khanday, S. S. Bandaru, R. C. Thomas, R. Y. Broadhurst, A. Venner, W. D. Todd, P. M. Fuller, E. Arrigoni, C. B. Saper, Role of serotonergic dorsal raphe neurons in hypercapnia-induced arousals. Nat Commun 11, 2769 (2020).

9. F. A. Mir, S. K. Jha, Locus Coeruleus Acid-Sensing Ion Channels Modulate Sleep-Wakefulness and State Transition from NREM to REM Sleep in the Rat. Neurosci Bull 37, 684–700 (2021).

10. M. Iwamoto, S. Yonekura, N. Atsumi, S. Hirabayashi, H. Kanazawa, Y. Kuniyoshi, Respiratory entrainment of the locus coeruleus modulates arousal level to avoid physical risks from external vibration. Sci Rep 13, 7069 (2023).

11. C. R. Sobrinho, B. M. Milla, J. Soto-Perez, T. S. Moreira, D. K. Mulkey, Histamine/H1 receptor signaling in the parafacial region increases activity of chemosensitive neurons and respiratory activity in rats. J Neurophysiol 128, 218–228 (2022).

12. P. L. Johnson, R. Moratalla, S. L. Lightman, C. A. Lowry, Are tuberomammillary histaminergic neurons involved in CO2-mediated arousal? Exp Neurol 193, 228–233 (2005).

13. Y. A. Li, J. Yao, X. Li, K. H. Hu, Arousal-promoting effect of the parabrachial nucleus and the underlying mechanisms: Recent advances. Prog Neuropsychopharmacol Biol Psychiatry 136, 111226 (2024).

14. J. T. McKenna, S. Thankachan, D. S. Uygun, C. Shukla, J. M. McNally, F. L. Schiffino, J. Cordeira, F. Katsuki, J. C. Zant, M. C. Gamble, K. Deisseroth, R. W. McCarley, R. E. Brown, R. E. Strecker, R. Basheer, Basal Forebrain Parvalbumin Neurons Mediate Arousals from Sleep Induced by Hypercarbia or Auditory Stimuli. Curr Biol 30, 2379–2385 e2374 (2020).

15. H. R. Smith, N. K. Leibold, D. A. Rappoport, C. M. Ginapp, B. S. Purnell, N. M. Bode, S. L. Alberico, Y. C. Kim, E. Audero, C. T. Gross, G. F. Buchanan, Dorsal Raphe Serotonin Neurons Mediate CO(2)-Induced Arousal from Sleep. J Neurosci 38, 1915–1925 (2018).

16. G. F. Buchanan, G. B. Richerson, Central serotonin neurons are required for arousal to CO2. Proc Natl Acad Sci U S A 107, 16354–16359 (2010).

17. A. Eban-Rothschild, G. Rothschild, W. J. Giardino, J. R. Jones, L. de Lecea, VTA dopaminergic neurons regulate ethologically relevant sleep-wake behaviors. Nat Neurosci 19, 1356–1366 (2016).

18. Y. Oishi, Y. Suzuki, K. Takahashi, T. Yonezawa, T. Kanda, Y. Takata, Y. Cherasse, M. Lazarus, Activation of ventral tegmental area dopamine neurons produces wakefulness through dopamine D(2)-like receptors in mice. Brain Struct Funct 222, 2907–2915 (2017).

19. X. Yu, W. Li, Y. Ma, K. Tossell, J. J. Harris, E. C. Harding, W. Ba, G. Miracca, D. Wang, L. Li, J. Guo, M. Chen, Y. Li, R. Yustos, A. L. Vyssotski, D. Burdakov, Q. Yang, H. Dong, N. P. Franks, W. Wisden, GABA and glutamate neurons in the VTA regulate sleep and wakefulness. Nat Neurosci 22, 106–119 (2019).

20. Y. Wang, D. Wang, X. Zhang, H. Li, S. Wang, Y. He, G. Zhao, H. Dong, J. Li, Dorsal Raphe Serotonergic Neurons-Ventral Tegmental Area Neural Pathway Promotes Wake From Sleep. CNS Neurosci Ther 30, e70141 (2024).

21. F. Naganuma, D. Kroeger, S. S. Bandaru, G. Absi, J. C. Madara, R. Vetrivelan, Lateral hypothalamic neurotensin neurons promote arousal and hyperthermia. PLoS Biol 17, e3000172 (2019).

22. H. L. Woodworth, J. A. Brown, H. M. Batchelor, R. Bugescu, G. M. Leinninger, Determination of neurotensin projections to the ventral tegmental area in mice. Neuropeptides 68, 57–74 (2018).

23. S. A. Oakman, P. L. Faris, P. E. Kerr, C. Cozzari, B. K. Hartman, Distribution of pontomesencephalic cholinergic neurons projecting to substantia nigra differs significantly from those projecting to ventral tegmental area. J Neurosci 15, 5859–5869 (1995).

24. J. R. Cho, J. B. Treweek, J. E. Robinson, C. Xiao, L. R. Bremner, A. Greenbaum, V. Gradinaru, Dorsal Raphe Dopamine Neurons Modulate Arousal and Promote Wakefulness by Salient Stimuli. Neuron 94, 1205–1219 e1208 (2017).

25. R. T. Huckstepp, R. id Bihi, R. Eason, K. M. Spyer, N. Dicke, K. Willecke, N. Marina, A. V. Gourine, N. Dale, Connexin hemichannel-mediated CO2-dependent release of ATP in the medulla oblongata contributes to central respiratory chemosensitivity. J Physiol 588, 3901–3920 (2010).

26. V. M. Dospinescu, S. Nijjar, F. Spanos, J. Cook, E. de Wolf, M. A. Biscotti, M. Gerdol, N. Dale, Structural determinants of CO(2)-sensitivity in the beta connexin family suggested by evolutionary analysis. Commun Biol 2, 331 (2019).

27. D. H. Brotherton, S. Nijjar, C. G. Savva, N. Dale, A. D. Cameron, Structures of wild-type and a constitutively closed mutant of connexin26 shed light on channel regulation by CO(2). Elife 13, 93686 (2024).

28. L. Meigh, S. A. Greenhalgh, T. L. Rodgers, M. J. Cann, D. I. Roper, N. Dale, CO(2)directly modulates connexin 26 by formation of carbamate bridges between subunits. Elife 2, e01213 (2013).

29. J. van de Wiel, L. Meigh, A. Bhandare, J. Cook, S. Nijjar, R. Huckstepp, N. Dale, Connexin26 mediates CO(2)-dependent regulation of breathing via glial cells of the medulla oblongata. Commun Biol 3, 521 (2020).

30. J. I. Nagy, A. V. Ionescu, B. D. Lynn, J. E. Rash, Coupling of astrocyte connexins Cx26, Cx30, Cx43 to oligodendrocyte Cx29, Cx32, Cx47: Implications from normal and connexin32 knockout mice. Glia 44, 205-218 (2003).

31. M. Vandecasteele, J. Glowinski, L. Venance, Connexin mRNA expression in single dopaminergic neurons of substantia nigra pars compacta. Neurosci Res 56, 419–426 (2006).

32. M. B. Singh, J. A. White, E. J. McKimm, M. M. Milosevic, S. D. Antic, Mechanisms of Spontaneous Electrical Activity in the Developing Cerebral Cortex-Mouse Subplate Zone. Cereb Cortex 29, 3363–3379 (2019).

33. A. R. Moore, W. L. Zhou, C. L. Sirois, G. S. Belinsky, N. Zecevic, S. D. Antic, Connexin hemichannels contribute to spontaneous electrical activity in the human fetal cortex. Proc Natl Acad Sci U S A 111, E3919–3928 (2014).

34. E. Hill, N. Dale, M. J. Wall, Moderate Changes in CO(2) Modulate the Firing of Neurons in the VTA and Substantia Nigra. iScience 23, 101343 (2020).

35. S. Nijjar, D. Maddison, L. Meigh, E. de Wolf, T. Rodgers, M. J. Cann, N. Dale, Opposing modulation of Cx26 gap junctions and hemichannels by CO(2). J Physiol 599, 103–118 (2021).

36. S. K. Ogawa, J. Y. Cohen, D. Hwang, N. Uchida, M. Watabe-Uchida, Organization of monosynaptic inputs to the serotonin and dopamine neuromodulatory systems. Cell Rep 8, 1105–1118 (2014).

37. W. J. Zou, Y. L. Song, M. Y. Wu, X. T. Chen, Q. L. You, Q. Yang, Z. Y. Luo, L. Huang, Y. Kong, J. Feng, D. X. Fang, X. W. Li, J. M. Yang, L. Mei, T. M. Gao, A discrete serotonergic circuit regulates vulnerability to social stress. Nat Commun 11, 4218 (2020).

38. H. L. Wang, S. Zhang, J. Qi, H. Wang, R. Cachope, C. A. Mejias-Aponte, J. A. Gomez, G. E. Mateo-Semidey, G. M. J. Beaudoin, C. A. Paladini, J. F. Cheer, M. Morales, Dorsal Raphe Dual Serotonin-Glutamate Neurons Drive Reward by Establishing Excitatory Synapses on VTA Mesoaccumbens Dopamine Neurons. Cell Rep 26, 1128–1142 e1127 (2019).

39. A. Belmer, P. M. Klenowski, O. L. Patkar, S. E. Bartlett, Mapping the connectivity of serotonin transporter immunoreactive axons to excitatory and inhibitory neurochemical synapses in the mouse limbic brain. Brain Struct Funct 222, 1297–1314 (2017).

40. R. Musa-Aziz, L. M. Chen, M. F. Pelletier, W. F. Boron, Relative CO2/NH3 selectivities of AQP1, AQP4, AQP5, AmtB, and RhAG. Proc Natl Acad Sci U S A 106, 5406-5411 (2009).

41. M. Alishahi, R. Kamali, A novel molecular dynamics study of CO(2) permeation through aquaporin-5. Eur Phys J E Soft Matter 42, 151 (2019).

42. J. E. Contreras, H. A. Sanchez, E. A. Eugenin, D. Speidel, M. Theis, K. Willecke, F. F. Bukauskas, M. V. Bennett, J. C. Saez, Metabolic inhibition induces opening of unapposed connexin 43 gap junction hemichannels and reduces gap junctional communication in cortical astrocytes in culture. Proc Natl Acad Sci U S A 99, 495–500 (2002).

43. Z. C. Ye, M. S. Wyeth, S. Baltan-Tekkok, B. R. Ransom, Functional hemichannels in astrocytes: a novel mechanism of glutamate release. J Neurosci 23, 3588–3596 (2003).

44. S. Faton, J. P. Tassin, F. Duranton, D. Bagnol, A. D. Lajoix, 5-HT(2C) receptors in the ventral tegmental area, but not in the arcuate nucleus, mediate the hypophagic and hypolocomotor effects of the selective 5-HT(2C) receptor agonist AR231630 in rats. Behav Brain Res 347, 234–241 (2018).

45. D. V. Herin, M. J. Bubar, P. K. Seitz, M. L. Thomas, G. R. Hillman, Y. I. Tarasenko, P. Wu, K. A. Cunningham, Elevated Expression of Serotonin 5-HT(2A) Receptors in the Rat Ventral Tegmental Area Enhances Vulnerability to the Behavioral Effects of Cocaine. Front Psychiatry 4, 2 (2013).

46. M. J. Bubar, K. A. Cunningham, Distribution of serotonin 5-HT2C receptors in the ventral tegmental area. Neuroscience 146, 286–297 (2007).

47. Z. A. Rodd, R. L. Bell, S. M. Oster, J. E. Toalston, T. J. Pommer, W. J. McBride, J. M. Murphy, Serotonin-3 receptors in the posterior ventral tegmental area regulate ethanol self-administration of alcohol-preferring (P) rats. Alcohol 44, 245–255 (2010).

48. Z. A. Rodd, V. E. Gryszowka, J. E. Toalston, S. M. Oster, D. Ji, R. L. Bell, W. J. McBride, The reinforcing actions of a serotonin-3 receptor agonist within the ventral tegmental area: evidence for subregional and genetic differences and involvement of dopamine neurons. J Pharmacol Exp Ther 321, 1003–1012 (2007).

49. Y. Minabe, C. R. Ashby, Jr., J. E. Schwartz, R. Y. Wang, The 5-HT3 receptor antagonists LY 277359 and granisetron potentiate the suppressant action of apomorphine on the basal firing rate of ventral tegmental dopamine cells. Eur J Pharmacol 209, 143–150 (1991).

50. Z. Liu, E. B. Bunney, S. B. Appel, M. S. Brodie, Serotonin reduces the hyperpolarization-activated current (Ih) in ventral tegmental area dopamine neurons: involvement of 5-HT2 receptors and protein kinase C. J Neurophysiol 90, 3201–3212 (2003).

51. M. S. Brodie, E. B. Bunney, Serotonin potentiates dopamine inhibition of ventral tegmental area neurons in vitro. J Neurophysiol 76, 2077–2082 (1996).

52. T. Okamoto, M. T. Harnett, H. Morikawa, Hyperpolarization-activated cation current (Ih) is an ethanol target in midbrain dopamine neurons of mice. J Neurophysiol 95, 619–626 (2006).

53. J. McDaid, M. A. McElvain, M. S. Brodie, Ethanol effects on dopaminergic ventral tegmental area neurons during block of Ih: involvement of barium-sensitive potassium currents. J Neurophysiol 100, 1202–1210 (2008).

54. P. Zhong, C. R. Vickstrom, X. Liu, Y. Hu, L. Yu, H. G. Yu, Q. S. Liu, HCN2 channels in the ventral tegmental area regulate behavioral responses to chronic stress. Elife 7, e32684 (2018).

55. L. Mu, X. Liu, H. Yu, C. R. Vickstrom, V. Friedman, T. J. Kelly, Y. Hu, W. Su, S. Liu, J. R. Mantsch, Q. S. Liu, cAMP-mediated upregulation of HCN channels in VTA dopamine neurons promotes cocaine reinforcement. Mol Psychiatry 28, 3930–3942 (2023).

56. S. H. Sheu, S. Upadhyayula, V. Dupuy, S. Pang, F. Deng, J. Wan, D. Walpita, H. A. Pasolli, J. Houser, S. Sanchez-Martinez, S. E. Brauchi, S. Banala, M. Freeman, C. S. Xu, T. Kirchhausen, H. F. Hess, L. Lavis, Y. Li, S. Chaumont-Dubel, D. E. Clapham, A serotonergic axon-cilium synapse drives nuclear signaling to alter chromatin accessibility. Cell 185, 3390–3407 e3318 (2022).

57. M. C. Fiori, V. Figueroa, M. E. Zoghbi, J. C. Saez, L. Reuss, G. A. Altenberg, Permeation of calcium through purified connexin 26 hemichannels. J Biol Chem 287, 40826–40834 (2012).

58. W. Lopez, J. Ramachandran, A. Alsamarah, Y. Luo, A. L. Harris, J. E. Contreras, Mechanism of gating by calcium in connexin hemichannels. Proc Natl Acad Sci U S A 113, E7986–E7995 (2016).

59. I. Delvendahl, L. Jablonski, C. Baade, V. Matveev, E. Neher, S. Hallermann, Reduced endogenous Ca2+ buffering speeds active zone Ca2+ signaling. Proc Natl Acad Sci U S A 112, E3075–3084 (2015).

60. V. Beaumont, A. Llobet, L. Lagnado, Expansion of calcium microdomains regulates fast exocytosis at a ribbon synapse. Proc Natl Acad Sci U S A 102, 10700–10705 (2005).

61. N. Dale, J. Butler, V. M. Dospinescu, S. Nijjar, Channel-mediated ATP release in the nervous system. Neuropharmacology 227, 109435 (2023).

62. Z. Ma, A. Taruno, M. Ohmoto, M. Jyotaki, J. C. Lim, H. Miyazaki, N. Niisato, Y. Marunaka, R. J. Lee, H. Hoff, R. Payne, A. Demuro, I. Parker, C. H. Mitchell, J. Henao-Mejia, J. E. Tanis, I. Matsumoto, M. G. Tordoff, J. K. Foskett, CALHM3 Is Essential for Rapid Ion Channel-Mediated Purinergic Neurotransmission of GPCR-Mediated Tastes. Neuron 98, 547–561 e510 (2018).

63. A. Taruno, K. Nomura, T. Kusakizako, Z. Ma, O. Nureki, J. K. Foskett, Taste transduction and channel synapses in taste buds. Pflugers Arch 473, 3–13 (2021).

64. R. A. Romanov, R. S. Lasher, B. High, L. E. Savidge, A. Lawson, O. A. Rogachevskaja, H. Zhao, V. V. Rogachevsky, M. F. Bystrova, G. D. Churbanov, I. Adameyko, T. Harkany, R. Yang, G. J. Kidd, P. Marambaud, J. C. Kinnamon, S. S. Kolesnikov, T. E. Finger, Chemical synapses without synaptic vesicles: Purinergic neurotransmission through a CALHM1 channel-mitochondrial signaling complex. Sci Signal 11, eaao1815 (2018).

65. S. Soma, N. Hayatsu, K. Nomura, M. W. Sherwood, T. Murakami, Y. Sugiyama, N. Suematsu, T. Aoki, Y. Yamada, M. Asayama, M. Kaneko, K. Ohbayashi, M. Arizono, M. Ohtsuka, S. Hamada, I. Matsumoto, Y. Iwasaki, N. Ohno, Y. Okazaki, A. Taruno, Channel synapse mediates neurotransmission of airway protective chemoreflexes. Cell 188, 2687–2704 e2629 (2025).

66. L. Valencia-Torres, C. M. Olarte-Sanchez, D. J. Lyons, T. Georgescu, M. Greenwald-Yarnell, M. G. Myers, Jr., C. M. Bradshaw, L. K. Heisler, Activation of Ventral Tegmental Area 5-HT(2C) Receptors Reduces Incentive Motivation. Neuropsychopharmacology 42, 1511–1521 (2017).

67. C. Nocjar, B. L. Roth, E. A. Pehek, Localization of 5-HT(2A) receptors on dopamine cells in subnuclei of the midbrain A10 cell group. Neuroscience 111, 163–176 (2002).

68. E. Dale, A. L. Pehrson, T. Jeyarajah, Y. Li, S. C. Leiser, G. Smagin, C. K. Olsen, C. Sanchez, Effects of serotonin in the hippocampus: how SSRIs and multimodal antidepressants might regulate pyramidal cell function. CNS Spectr 21, 143–161 (2016).

69. M. K. Holt, N. Valderrama, M. J. Polanco, I. Hayter, E. G. Badenoch, S. Trapp, L. Rinaman, Modulation of stress-related behaviour by preproglucagon neurons and hypothalamic projections to the nucleus of the solitary tract. Mol Metab 91, 102076 (2025).

70. A. M. Bhandare, N. Dale, Neural correlate of reduced respiratory chemosensitivity during chronic epilepsy. Front Cell Neurosci 17, 1288600 (2023).

