## Supplementary material for "CO_2_-sensitive Cx26 hemichannels in dorsal raphe-ventral tegmental area pathway mediate hypercapnic arousal"

Lumei Huang et al.

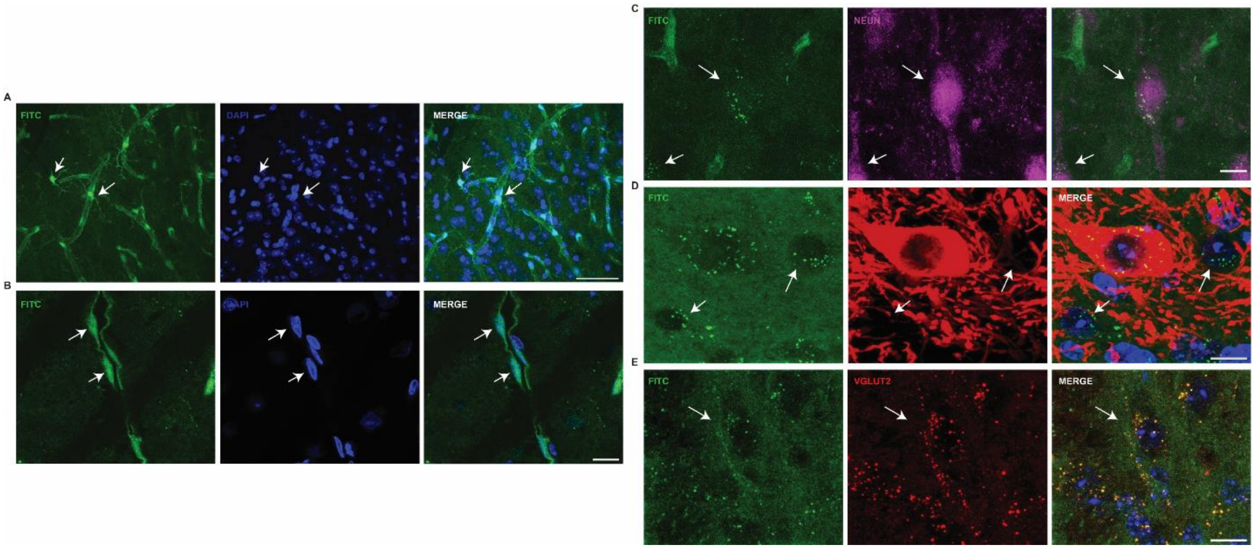


**Fig. S1. FITC labelled blood vessels and non-dopaminergic neurons in the VTA. (A)** Example confocal images showing FITC (green) labelled blood vessels and astrocytes around blood vessels (white arrow). Scale bar 10 µm. **(B)** Confocal images showing FITC (green) labelled endothelial cells. Scale bar 20 µm. **(C)** FITC-loaded puncta (green) clustered around NEUN positive cells (magenta) and non-dopaminergic neurons **(D)** in the VTA. E) FITC-loaded puncta colocalized with the excitatory presynaptic marker VGLUT2 (red). Scale bar for **C-E** 10 µm.

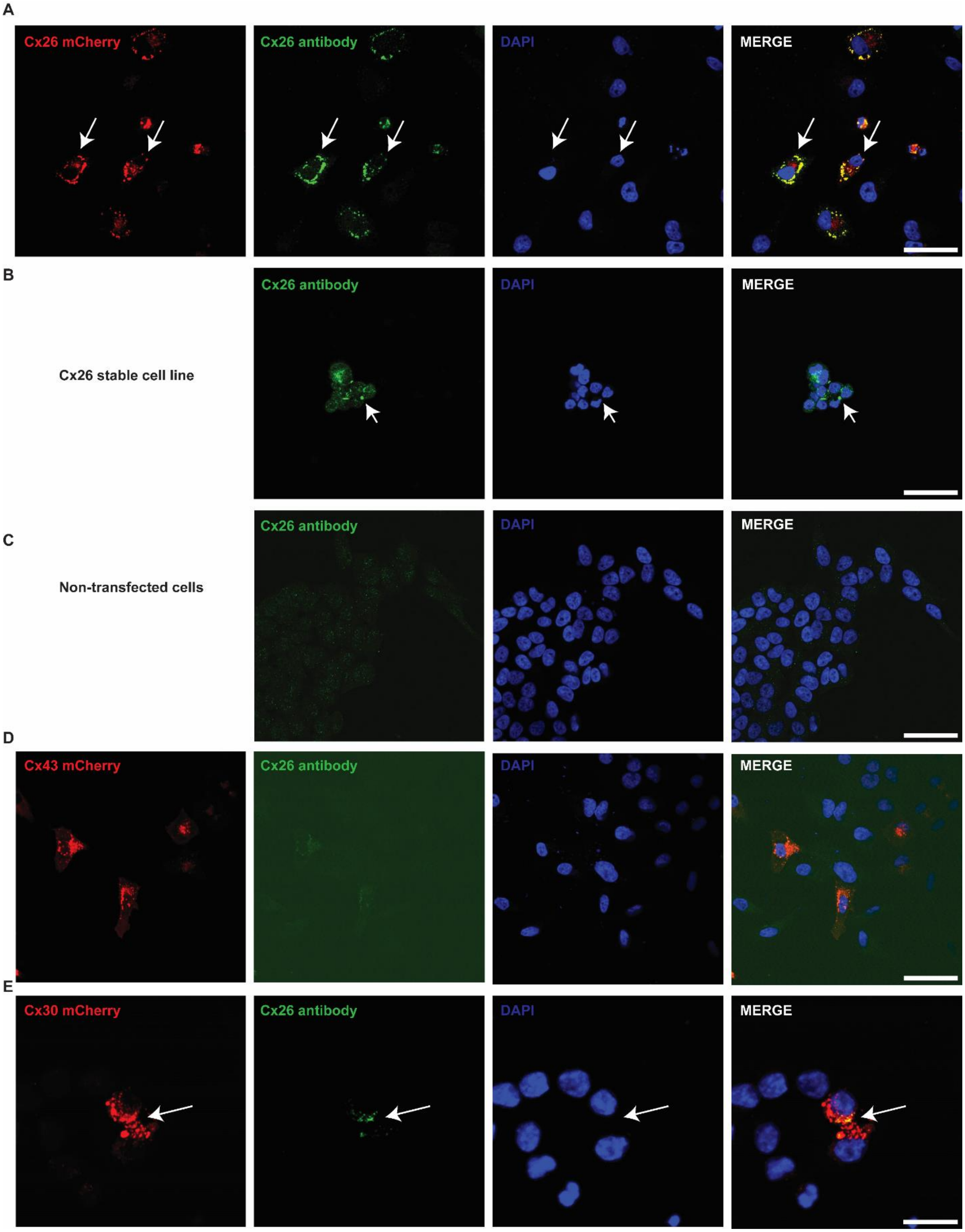
A
Cx26 mCherry
Cx26 antibody
DAPI
MERGE
B
Cx26 stable cell line
Cx26 antibody
DAPI
MERGE
C
Non-transfected cells
Cx26 antibody
DAPI
MERGE
D
Cx43 mCherry
Cx26 antibody
DAPI
MERGE
E
Cx30 mCherry
Cx26 antibody
DAPI
MERGE

**Fig. S2. Cx26 antibody validation in cultured cells.** (A) HeLa cells transfected with Cx26 mCherry stained of mouse monoclonal Cx26 antibody (green). Cx26 antibody colocalized with mCherry. (B) Mouse monoclonal Cx26 antibody (green) colocalized with Cx26 stable cell lines. (C) Mouse monoclonal Cx26 antibody (green) did not recognize non-transfected Hella cells. (D) Mouse monoclonal Cx26 antibody (green) did not recognize another $CO_2$-sensitive hemichannel Cx43 (mCherry). (E) Mouse monoclonal Cx26 antibody (green) weakly localises with the $CO_2$-sensitive hemichannel Cx30 (mCherry). A-D, scale bar 50 µm; E, scale bar 20 µm. The white arrows indicate colocalization.

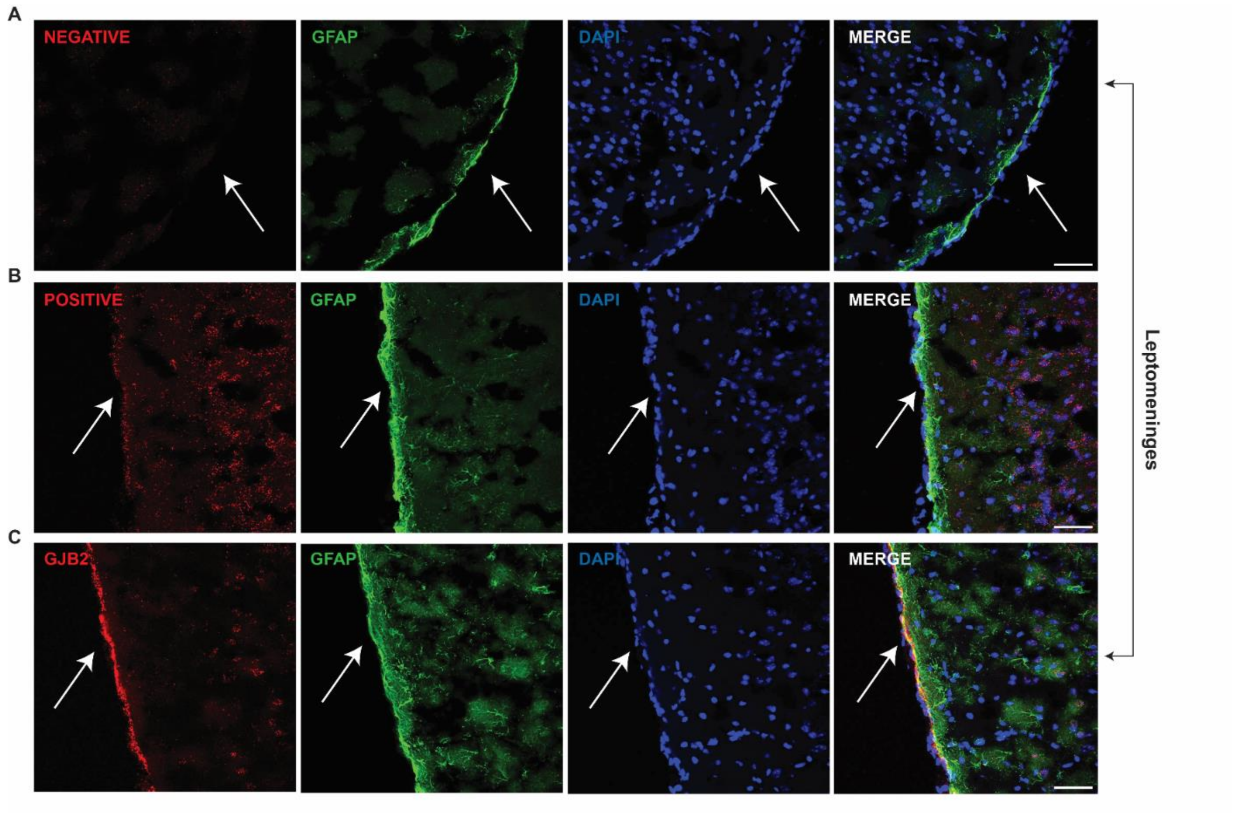


**Fig. S3. The expression of negative, positive and Gjb2 probes in leptomeninges. (A)** Negative probe (red) did not express on slices which were labelled with astrocyte maker (GFAP: green). (**B)** Positive probe (red) showed the housekeeping gene expressed on slices which were stained with astrocyte maker (GFAP: green), but only subtly expressed in the leptomeninges. (**C)** GJB2 (Red) densely expressed on leptomeninges. White arrows indicate the leptomeninges. Scale bar 50 μm.

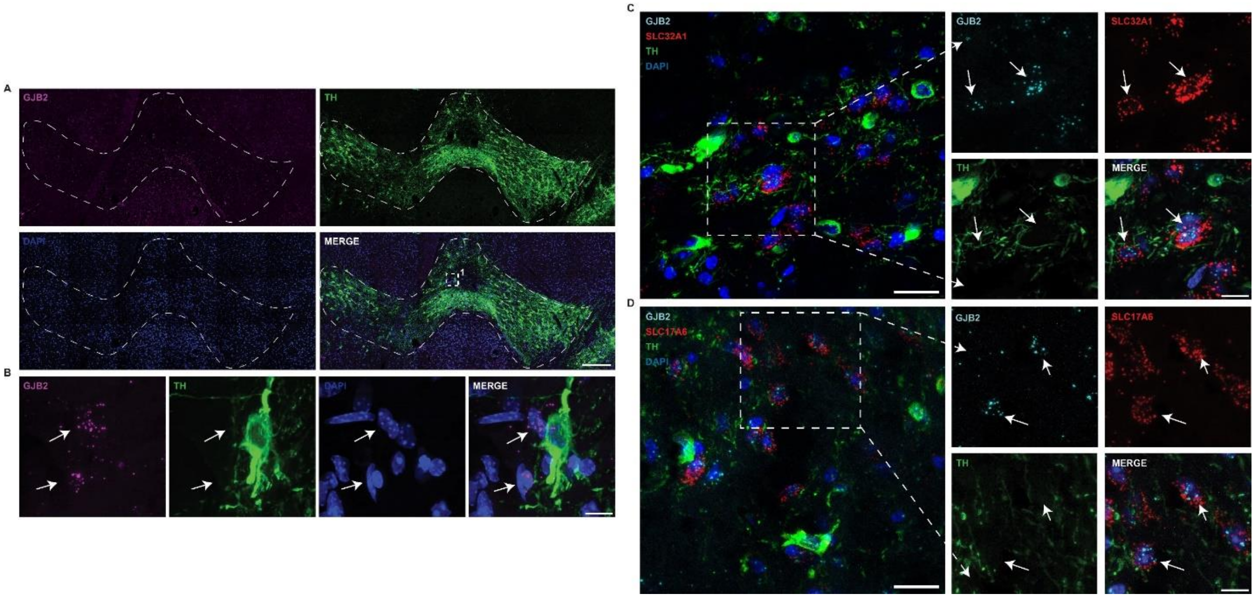


**Fig. S4. The detection of *Gjb2* mRNA in the VTA neurons.** RNAscope labelling combined with immunofluorescence was used to detect the *GJB2* expression in the VTA (TH staining)**. (A)** Tile images showing *GJB2* mRNA in the VTA labelled with TH (green) (Scale bar 200 µm). (**B)** Magnification of the white dashed areas in the tile images shows the expression of *GJB2* mRNA in non-TH negative cells, including both VGAT (Gene: SLC32A1) and VGLUT2 (gene: SLC17A6) in **C and D. (C)** Left panel showed merged images from GJB2 (cyan), SLC32A1 (red), TH (green) and DAPI (Blue). Right panels showed the high magnification images from the white square. GJB2 colocalized with some SLC32A1 (red) positive neurons. (**D)** Left panel showed merged images from GJB2 (cyan), SLC17A6 (red, VGLUT), TH (green) and DAPI (Blue). Right panels showed the high magnification images from the white square. GJB2 colocalized with some SLC17A6 (red, VGLUT2) positive neurons. White arrows indicate the colocalization. Scale for **C** and **D** left panels 20 µm, for right panels 10 µm.

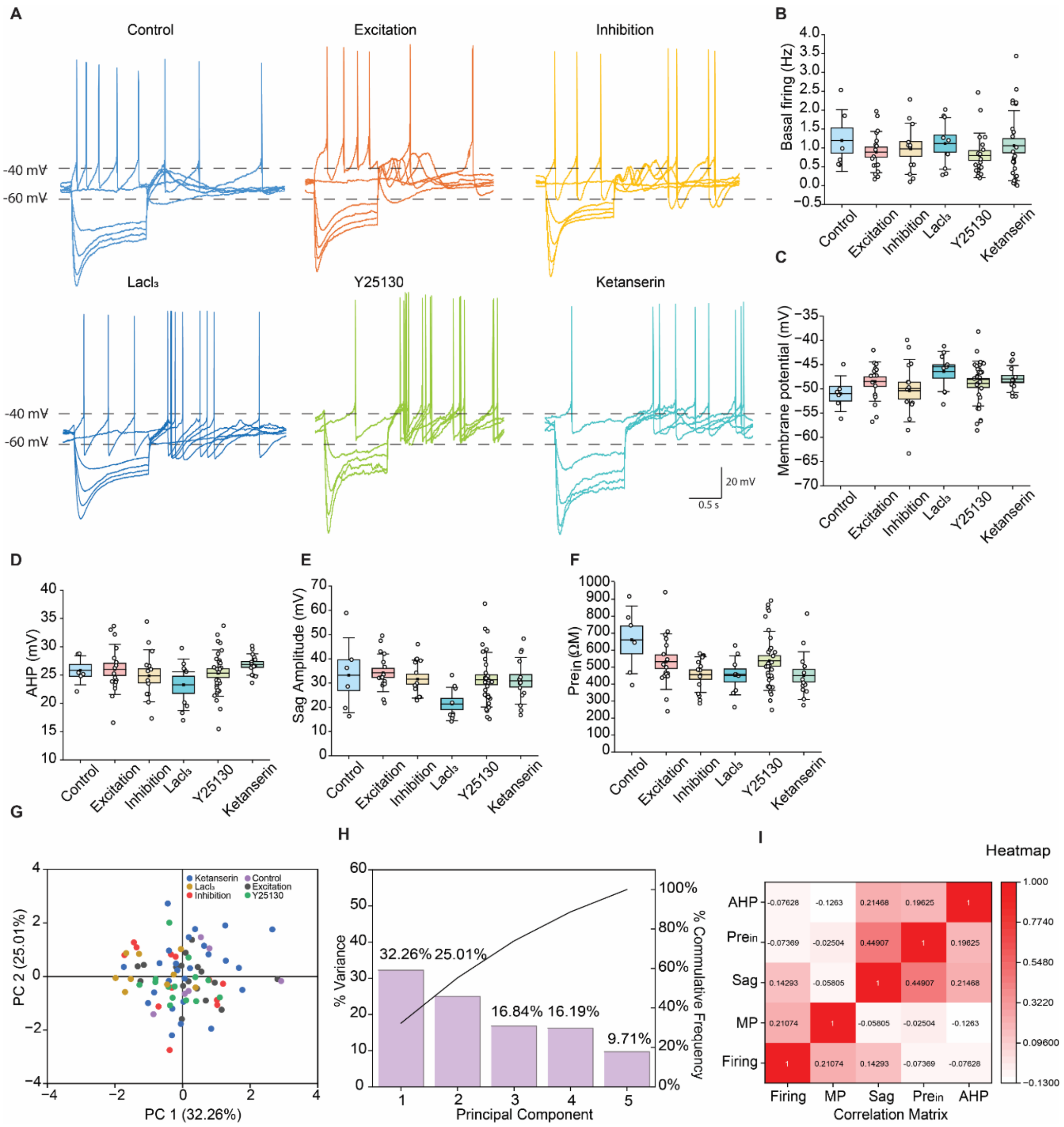


**Fig. S5. There are no significant differences in the electrophysiological properties of the VTA dopaminergic neurons across the groups. (A)** Voltage traces in response to current steps recorded from a dopamine neuron in each of the experimental groups (control, excited by $CO_2$, inhibited by $CO_2$, $LaCl_3$ treated, Y25130 treated and ketanserin treated). **B-F)** Summary data for electrophysiological properties: basal firing rate, membrane potential, AHP amplitude, sag amplitude, and pre-sag input resistance from the experimental groups. There are no significant differences in the measured parameters across the groups. Error bars are SEMs. (**G-I)** Principal component analysis (PCA) on the electrophysiological data from different groups **(B-F)**. (**G)** Comparison of data with reference to the first 2 principal components (PCs): PC1 vs PC2. (**H)** The graph shows the percentage of variance explained by each principal component (PC) (bars) and the cumulative percentage of variance (line). (**I)** shows the pair-wise estimated correlations obtained with 5 PCs. AHP: After hyperpolarization.

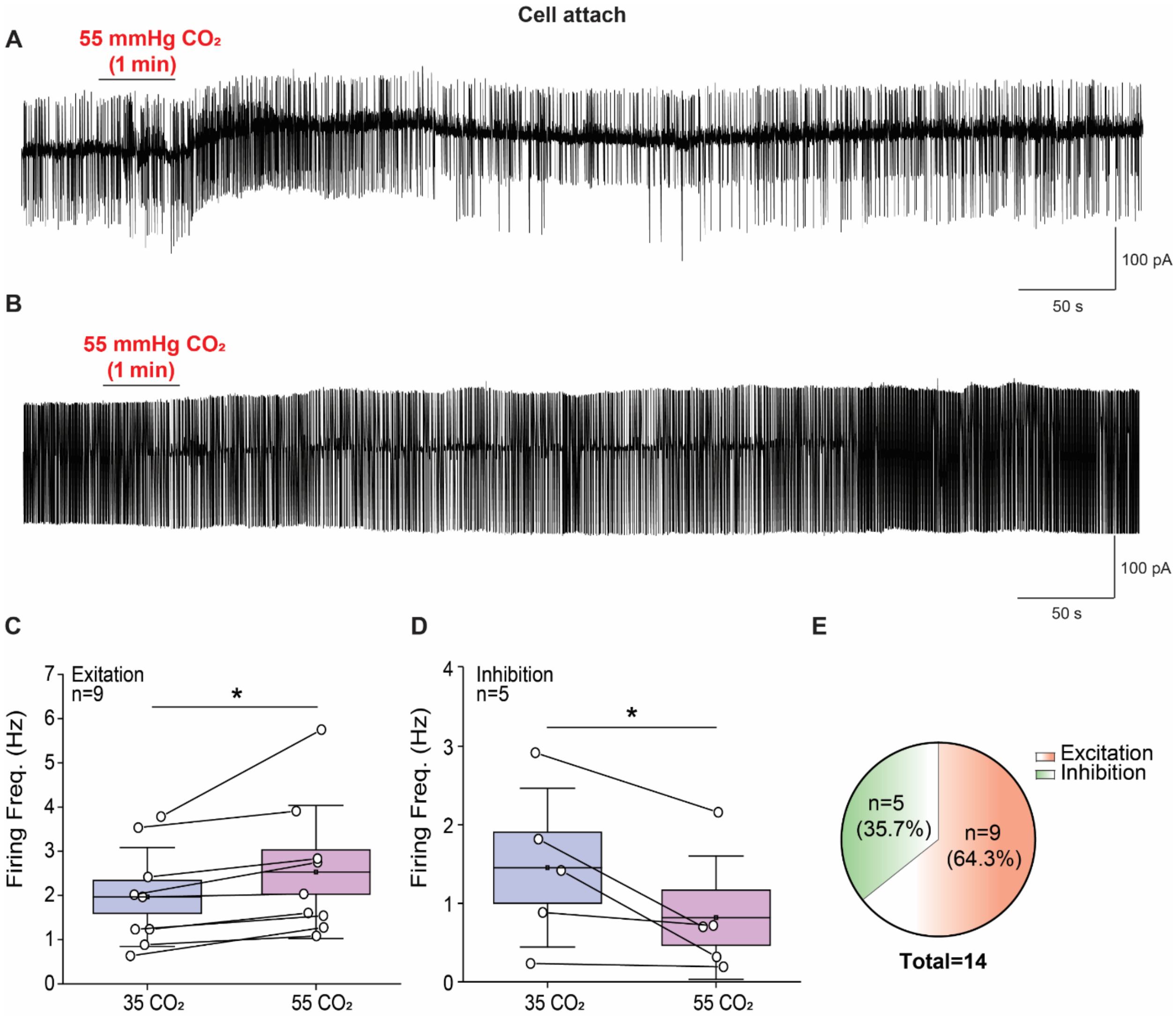


**Fig. S6. The effects of increased $PCO_2$ on VTA DA neuron firing rate in cell-attached patch clamp**. (**A and B)** Current traces from DA neurons showing spontaneous action potential firing in response to increased $PCO_2$ (1 minute from 35 to 55 mmHg). **(C and D)** Summary data illustrating bidirectional effects of elevated $PCO_2$ on VTA DA neuron firing rate. Elevated $PCO_2$ increased firing in one population of neurons (C) and decreased firing in another (D). Increased firing rate: n = 9 cells, paired t-test, *P = 0.017; decreased firing rate: n = 5 cells, paired t-test, *P = 0.049. Error bars represent SEM. (**E)** Pie chart showing the overall percentage of cells that were either excited or inhibited by increases in $PCO_2$.

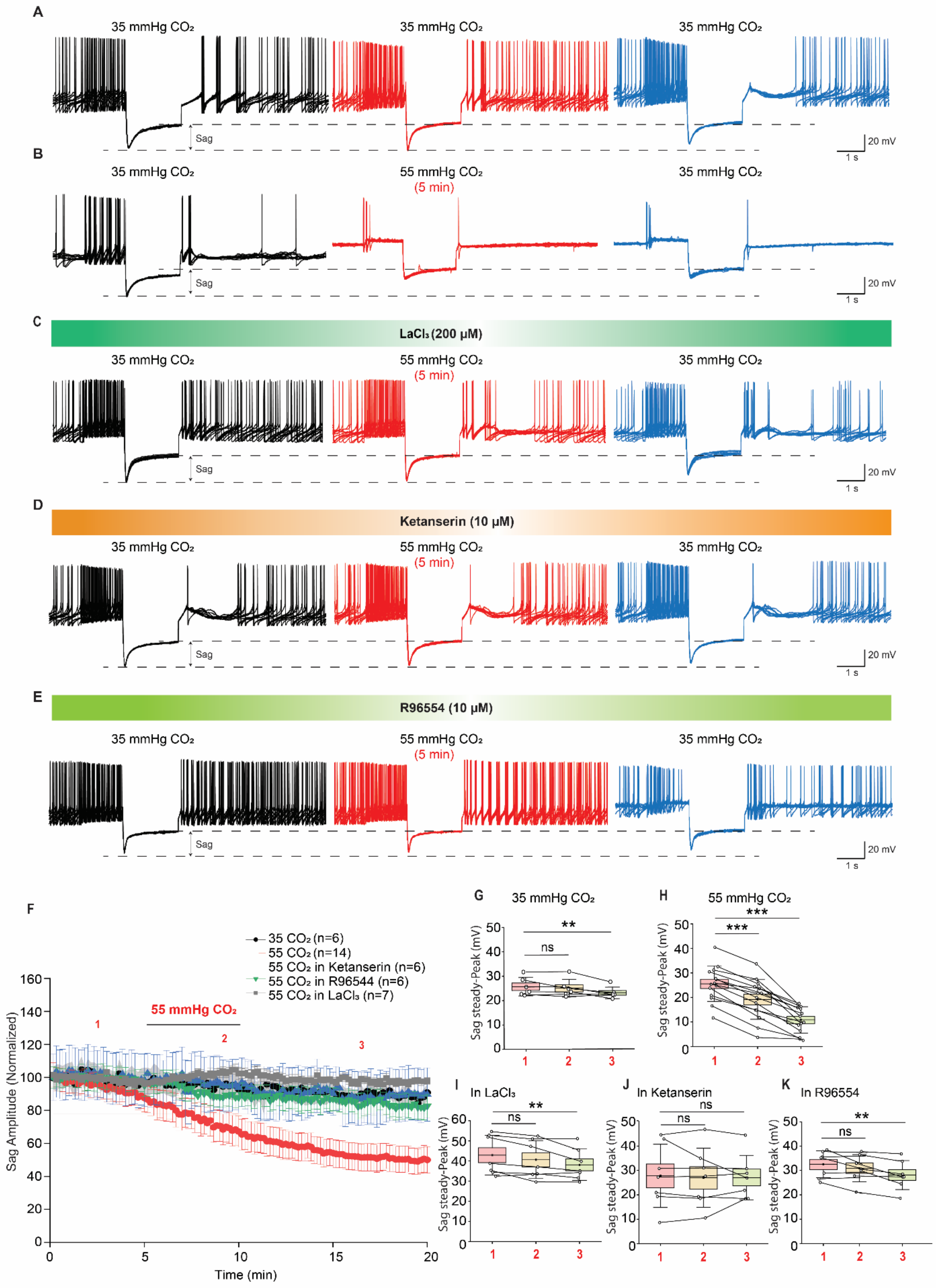
A
35 mmHg CO₂
35 mmHg CO₂
35 mmHg CO₂
Sag
20 mV
1 s
B
35 mmHg CO₂
55 mmHg CO₂
(5 min)
35 mmHg CO₂
Sag
20 mV
1 s
C
LaCl₃ (200 μM)
35 mmHg CO₂
55 mmHg CO₂
(5 min)
35 mmHg CO₂
Sag
20 mV
1 s
D
Ketanserin (10 μM)
35 mmHg CO₂
55 mmHg CO₂
(5 min)
35 mmHg CO₂
Sag
20 mV
1 s
E
R96554 (10 μM)
35 mmHg CO₂
55 mmHg CO₂
(5 min)
35 mmHg CO₂
Sag
20 mV
1 s
F
35 CO₂ (n=6)
55 CO₂ (n=14)
55 CO₂ in Ketanserin (n=6)
55 CO₂ in R96544 (n=6)
55 CO₂ in LaCl₃ (n=7)
55 mmHg CO₂
Sag Amplitude (Normalized)
Time (min)
G
35 mmHg CO₂
Sag steady-Peak (mV)
H
55 mmHg CO₂
Sag steady-Peak (mV)
I
In LaCl₃
Sag steady-Peak (mV)
J
In Ketanserin
Sag steady-Peak (mV)
K
In R96554
Sag steady-Peak (mV)
ns
**
***

**Fig. S7. Increased $PCO_2$ (55 mmHg $CO_2$ for 5 minutes) reduces the voltage sag amplitude of VTA DA neurons via $5HT_{2A}$ receptor activation.** Hyperpolarizing current steps were injected every 10 s to evoke the voltage sag. **(A–E)** Superimposed voltage traces (10 sweeps) illustrating the sag response to hyperpolarizing current injections. Sag amplitude was measured between the dashed lines (steady-state amplitude minus peak amplitude). **(F)** Time course of normalized sag amplitude for the different experimental groups. **(G–K)** Summary graphs showing changes in sag amplitude under the indicated experimental conditions. Mean sag amplitudes were calculated over 1-min periods at the time points indicated in **(F)**. **(G)** Sag amplitude remained relatively stable when $PCO_2$ was maintained at 35 mmHg. **(H)** Exposure to 55 mmHg $CO_2$ reduced sag amplitude. **(I)** The hemichannel blocker $LaCl_3$ abolished the reduction in sag amplitude. **(J)** The 5-$HT_2$ receptor antagonist ketanserin abolished the reduction in sag amplitude. **(K)** The selective 5-$HT_2A$ receptor antagonist R96554 abolished the reduction in sag amplitude. Data are mean ± SEM. Sag amplitudes were: **(G)** n = 6 cells; **(H)** n = 14 cells; **(I)** n = 7 cells; **(J)** n = 6 cells; **(K)** n = 6 cells). Statistical significance was assessed using one-way repeated-measures ANOVA followed by Tukey's multiple-comparisons test. Baseline versus 55 mmHg $CO_2$: (H) ***$P < 0.0001$, (I) $P = 0.25$, (J) $P = 0.92$, (K) $P = 0.22$.

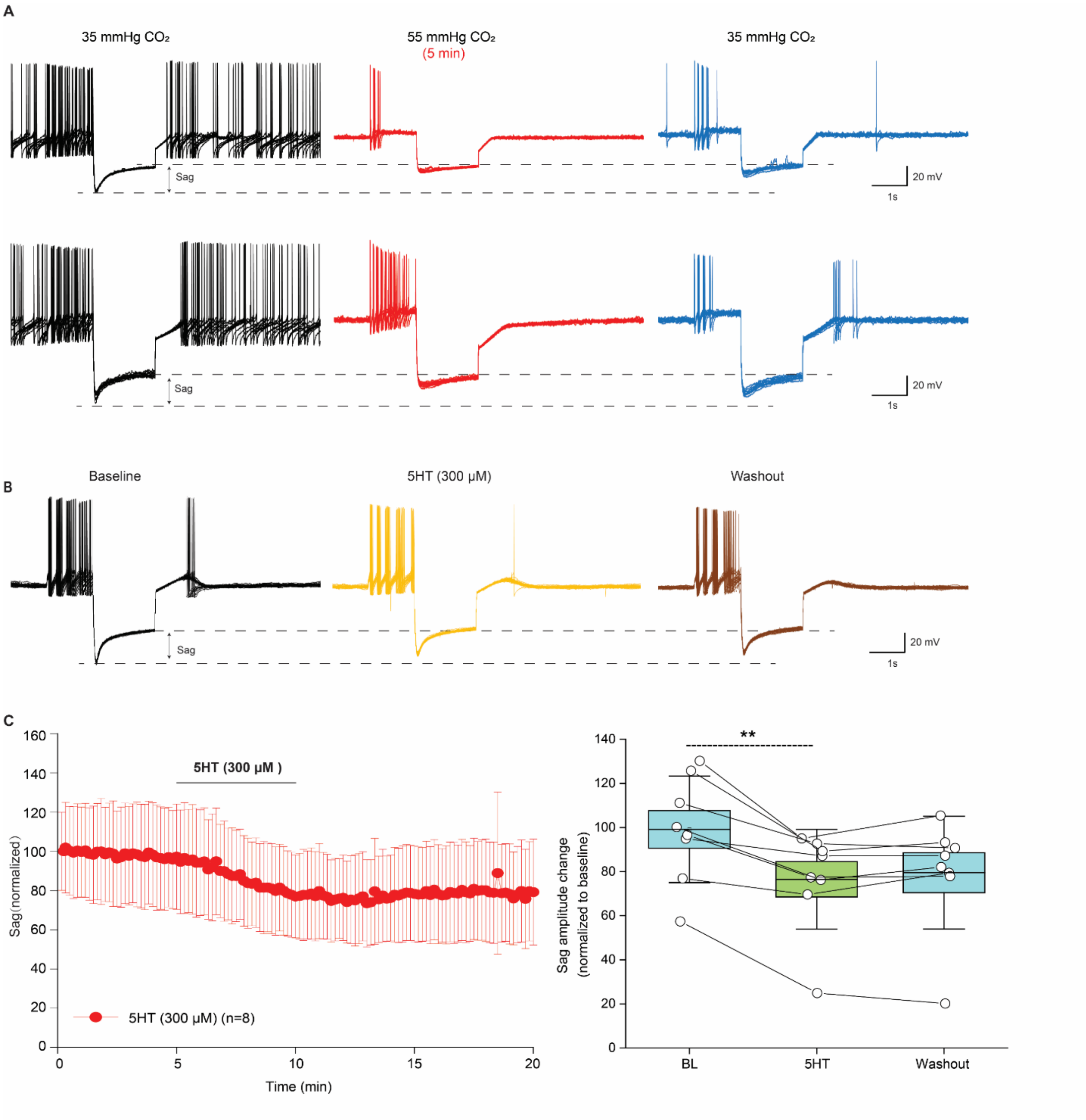


**Fig. S8. The inhibitory effects of raised PCO2 on VTA DA neuron hyperpolarising sag amplitude can be partially reversed by prolonged wash and can be mimicked by 5HT application. (A)** Two example recordings from DA VTA neurons showing that washing in 35 mm Hg $CO_2$ for ~ 40 minutes led to a partial recovery of sag amplitude that was inhibited by 55 mm Hg $CO_2$ (blue traces). (**B**) Application 5HT (300 µM) reduced sag amplitude. (**C**) Summary of the time course (Left) and quantification (Right) of 5HT effects on normalised sag amplitude ($n$ = 8). Data are mean ± SEM. Statistical significance was assessed using Friedman ANOVA followed by Dunn's multiple-comparisons test. **P = 0.003.

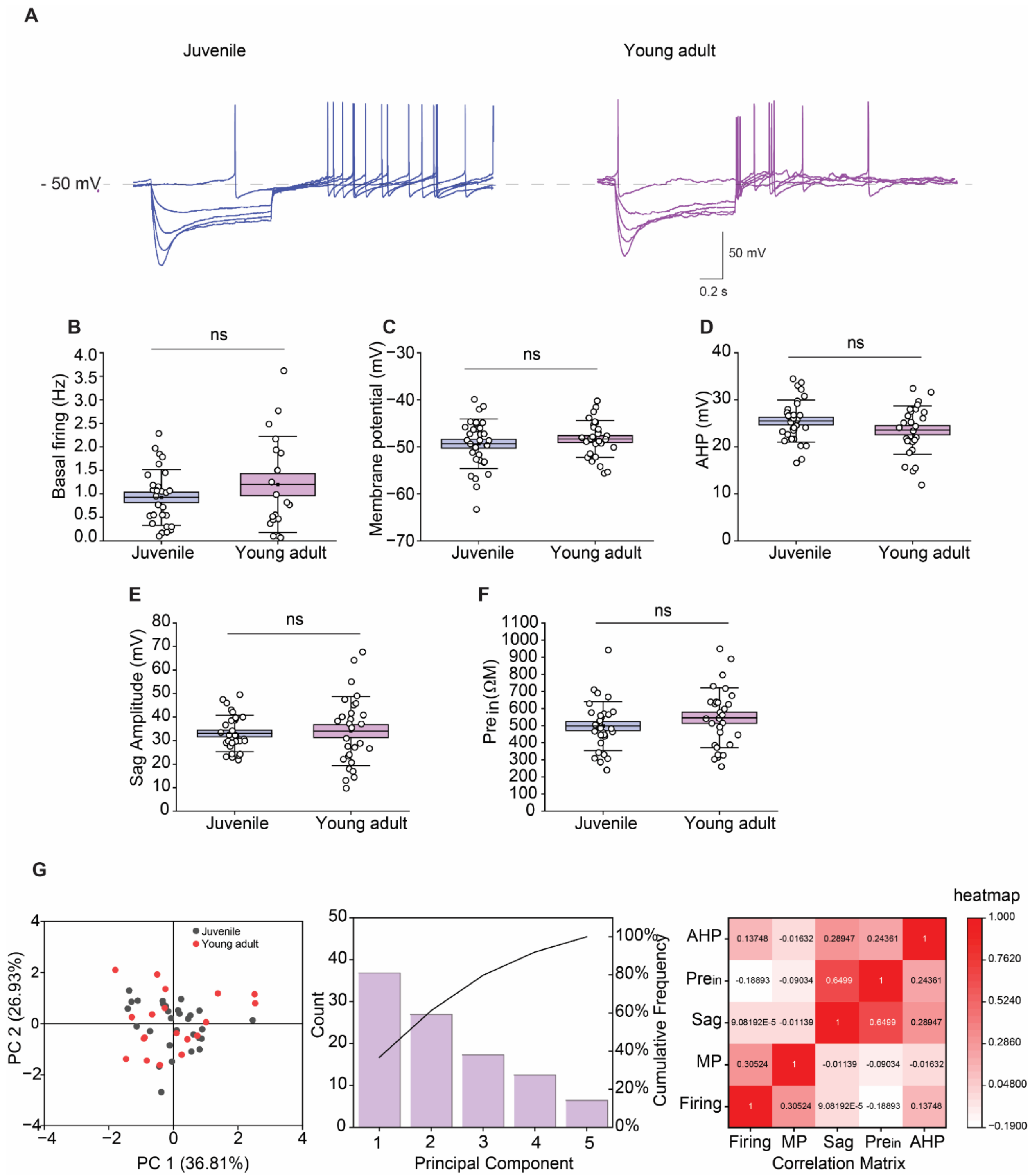


**Fig. S9. Electrophysiological characteristics of VTA DA neurons are similar in juvenile and young adult mice.** **(A)** Representative current-voltage traces recorded from juvenile (3–4 weeks) and young adult (3–4 months) VTA DA neurons. **(B)** Comparison of basal firing rate, membrane potential, AHP amplitude, sag amplitude, and pre-sag input resistance between juvenile and young adult mice. No significant differences were detected between groups. **(G–I)** Principal component analysis (PCA) of electrophysiological properties from juvenile and young adult VTA DA neurons. **(G)** Distribution of data according to the first two principal components (PC1 and PC2). **(H)** Percentage of variance explained by each principal component (bars) and cumulative variance explained (line). **(I)** Pairwise correlations estimated using the first five principal components. AHP, afterhyperpolarization. Data are mean ± SEM. Statistical significance was assessed using unpaired two-sample t-tests. Basal firing rate, P = 0.25; membrane potential, P = 0.41; AHP amplitude, P = 0.12; sag amplitude, P = 0.74; pre-sag input resistance, P = 0.24. ns, not significant.

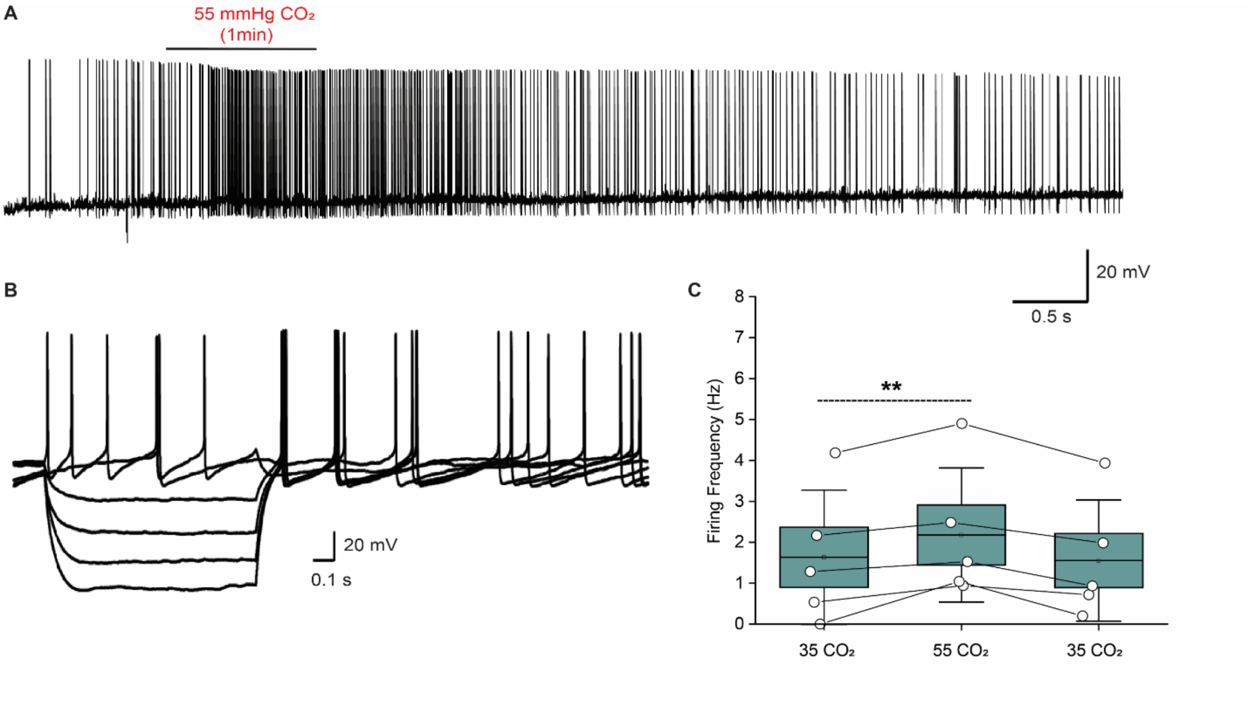


**Fig. S10. DR neurons respond to $CO_2$. (A)** 1 min 55 mmHg$CO_2$ increased the firing rate of DR neurons. (**B)** Representative current-voltage traces for DR neurons. (**C)** Summary of firing frequency at 35 and 55 mmHg $CO_2$. Compared to baseline (35 mmHg $CO_2$), 1 min 55 mmHg $CO_2$ significantly increased the firing frequency. Data are mean ± SEM. Statistical significance was assessed using one-way repeated-measures ANOVA followed by Tukey's multiple-comparisons test. **P = 0.008.

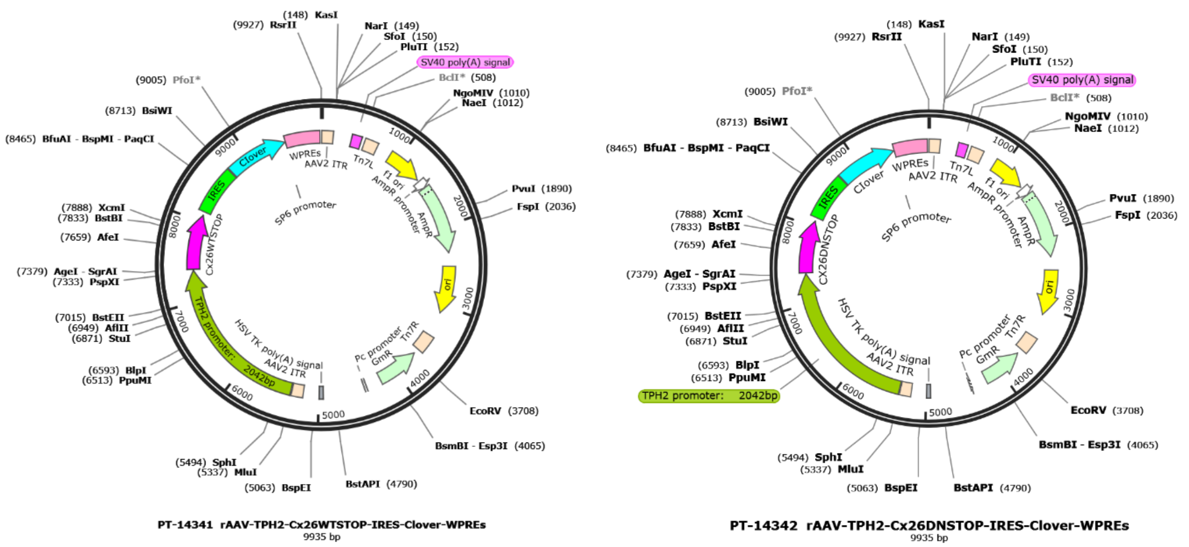


**Fig. S11.** Schematic representation of the recombinant AAV vector rAAV-TPH2-Cx26WT/DNSTOP-IRES-Clover-WPREs. The recombinant adeno-associated virus (rAAV) expression cassette consists of a 5′ AAV2 inverted terminal repeat (ITR), the mouse TPH2 promoter (2.0 kb), the coding sequence of wild-type connexin 26 (Cx26WT/DN) terminated by a stop codon, an internal ribosome entry site (IRES), the fluorescent reporter Clover, and the woodchuck hepatitis virus posttranscriptional regulatory element (WPREs), followed by a 3′ AAV2 ITR. The TPH2 promoter drives serotonergic neuron–specific expression of Cx26WT/DN, while the IRES enables independent translation of Clover from the same transcript for visualization of transduced cells. The construct was packaged into AAV particles for in vivo delivery.

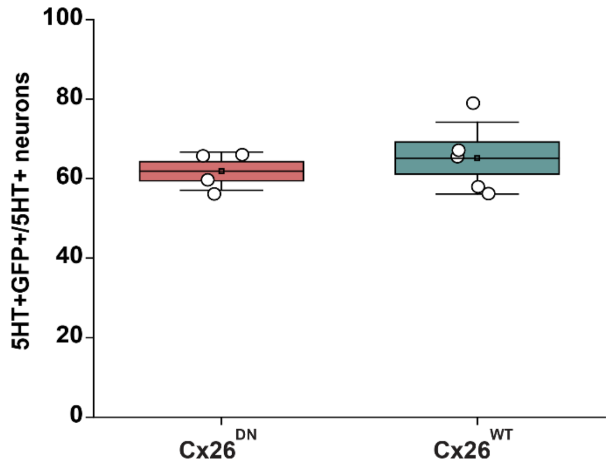


**Fig. S12. Virus infected proportion of DR 5HT neurons in Cx26$^{WT}$ and Cx26$^{ND}$ mice.** Cx26$^{WT}$ (4 animals): 62% ± 2.41 vs Cx26$^{DN}$ (5 animals): 65% ± 4.04. Data was presented as Mean ± SEM. *P*-value was determined by two sample t-test. P = 0.53.
